# Behavioral and fMRI evidence that arousal enhances bottom-up attention and memory selectivity in young but not older adults

**DOI:** 10.1101/2021.07.02.450802

**Authors:** Sara N. Gallant, Briana L. Kennedy, Shelby L. Bachman, Ringo Huang, Tae-Ho Lee, Mara Mather

## Abstract

During a challenge or emotional experience, increases in arousal help us focus on the most salient or relevant details and ignore distracting stimuli. The noradrenergic system integrates signals about arousal states throughout the brain and helps coordinate this adaptive attentional selectivity. However, age-related changes in the noradrenergic system and attention networks in the brain may reduce the efficiency of arousal to modulate selective processing in older adults. In the current neuroimaging study, we examined age differences in how arousal affects bottom-up attention to category-selective stimuli differing in perceptual salience. We found a dissociation in how arousal modulates selective processing in the young and older brain. In young adults, emotionally arousing sounds enhanced selective incidental memory and brain activity in the extrastriate body area for salient versus non-salient images of bodies. Older adults showed no such advantage in selective processing under arousal. These age differences could not be attributed to changes in the arousal response or less neural distinctiveness in old age. Rather, our results suggest that, relative to young adults, older adults become less effective at focusing on salient over non-salient details during increases in emotional arousal.

## 1. Introduction

Increases in arousal help tune attention to relevant aspects of our environment (Mather & Sutherland, 2011). This adaptive response can be triggered by spikes in arousal associated with a threat to survival or more common events like hearing a jarring sound, completing an effortful task, or experiencing certain emotions. The locus coeruleus (LC), the brain’s primary source of noradrenaline (NA), integrates signals about arousal states in the brain via its broad efferent network (Aston-Jones & Cohen, 2005; Samuels & Szabadi, 2008). LC neurons have both tonic and phasic modes of activity. The tonic mode is characterized by elevated baseline LC activity with low levels of phasic LC firing. The phasic mode, in contrast, is characterized by moderate baseline LC activity with high levels of phasic LC firing (Aston-Jones & Cohen, 2005). During the phasic activity mode, excitation of highly active neurons and suppression of less active neurons is enhanced, facilitating task-relevant behavior. As a result, during an arousing event, cognitive acuity may be improved (Sara, 2009; Sara & Bouret, 2012), enhancing attention and memory for salient information (Clewett et al., 2018; Jacobs et al., 2020; Lee et al., 2013, 2018). According to the glutamate amplifies noradrenergic effects (GANE) theory, this gain in selectivity under arousal is caused by high glutamate levels at the site of high priority representations, which interacts with local NA. These glutamate-NA interactions are thought to then enhance processing of high priority representations and laterally suppress processing of low priority representations (Mather et al., 2016).

Recent work suggests that aging may lead to a decline in the noradrenergic modulation of cognitive selectivity under arousal (Durbin et al., 2018; Gallant et al., 2020; Lee et al., 2018). For example, a tone conditioned to predict shock (CS+) selectively increased young adults’ brain activity for salient versus non-salient stimuli relative to a tone that did not predict shock (CS-; Lee et al., 2018). In contrast, the CS+ increased older adults’ brain activity for both salient and non-salient stimuli, suggesting that arousal enhanced their attention to distracting stimuli. One possibility is that these findings stem from age-related changes in the noradrenergic system (Mather & Harley, 2016; Robertson, 2013). The LC is the first place in the brain that Alzheimer’s related tau pathology is seen (Braak & Del Tredici, 2011). Moreover, as some LC neurons are damaged or lost with age, noradrenergic activity increases, potentially as a compensatory mechanism (Weinshenker, 2018). This may translate to higher levels of baseline LC activity that would limit the range of the arousal response in an older brain (Mather, 2020). As a result, functions associated with the LC-NA system, such as modulating cognition, could be disrupted. Consistent with this, LC structural integrity in later life has been linked with outcomes across cognitive domains such as verbal fluency and intelligence (Elman et al., 2021; Liu et al., 2020), learning and memory (Dahl et al., 2019), and emotional memory (Hämmerer et al., 2018).

Age-related declines in cognitive selectivity under arousal may also occur because of changes in the interaction between the LC and frontoparietal brain regions (Lee et al., 2018). Attention networks in frontoparietal regions contribute to selectivity by encoding the priority of information and guiding attention to these prioritized inputs (Ptak, 2012). A dorsal attention network selects stimuli based on internal goals and is centered on the intraparietal and superior frontal cortex. A ventral attention network detects salient and unexpected external stimuli and includes the ventral frontal cortex and temporoparietal junction, lateralized to the right hemisphere (Corbetta & Shulman, 2002). These frontoparietal regions are highly innervated by NA neurons (Samuels & Szabadi, 2008) and have been implicated in noradrenergic modulation of cognitive performance (Corbetta et al., 2008; Corbetta & Shulman, 2002; Sara & Bouret, 2012). Under arousal, the phasic LC response that amplifies neural gain (Aston-Jones & Cohen, 2005) is thought to modulate activity within frontoparietal regions, guiding detection of goal-relevant stimuli (Corbetta et al., 2008; Lee et al., 2018; Sara & Bouret, 2012). When comparing age groups, neuroimaging work has shown that heightened arousal leads to greater functional connectivity of the LC and dorsal frontoparietal network in young than in older adults (Lee et al., 2018). Thus, an age-related decline in LC modulation of frontoparietal activity may explain the age-related decline in arousal-induced cognitive selectivity.

While Lee et al. (2018) demonstrated age differences in the interaction of arousal and perceptual selectivity, the bottom-up and top-down influences on attention and brain activity could not be dissociated. In their task, participants detected the spatial location of salient cues, which involved a combination of orienting to external cues with an internal goal of detecting their spatial locations. Furthermore, whether the age difference in arousal-induced selectivity in perception extended to subsequent memory is unknown. Additional research that targets bottom-up attention and memory performance is thus required to answer questions about whether arousal has a different impact on these processes in young and older adults.

In the current study, we tested age differences in how arousal modulates brain activity associated with bottom-up attention and incidental memory for salient versus non-salient images. We predicted that, relative to young adults, arousal would be less effective at amplifying older adults’ selective brain activity and memory for images competing in perceptual salience. To test our hypothesis, we combined behavioral, pupillary, and functional magnetic resonance imaging (fMRI) methods. In an MRI scanner, young and older adults completed a dot-probe task for category-selective stimuli that differed in perceptual salience under arousing and non-arousing conditions. Before images were presented, a negative or neutral sound clip was played to manipulate arousal levels. Outside of the scanner, a surprise recognition task tested memory for all salient and non-salient images seen during the dot-probe task.

Use of category-selective stimuli during the dot-probe task allowed us to test whether the effects of arousal on selectivity generalize to distinct category-selective regions in the occipitotemporal cortex. On each trial, stimuli included a pair of body and scene images that differed in perceptual salience. These categories were selected based on evidence that the extrastriate body area (EBA) selectively responds to human bodies (Downing et al., 2001) and the parahippocampal place area (PPA) selectively responds to scene images (Epstein & Kanwisher, 1998). Prior studies investigated the impact of arousal on selectivity in the fusiform face area (FFA; Lee et al., 2013) and PPA (Clewett et al., 2018; Lee et al., 2013, 2018), but none have generalized arousal’s effect to the EBA. We therefore localized brain activity in response to body and scene images and compared visual encoding processes across arousal conditions.

As previously described, neuroimaging studies have shown that top-down attention is supported by dorsal attention network regions of the frontal and parietal lobes whereas bottom-up attention, mediated by stimulus salience, activates right-lateralized ventral regions including the temporoparietal junction (TPJ; Bowling et al., 2020; Corbetta et al., 2008; Shomstein, 2012). The right TPJ, near the intersection of the angular gyrus and superior parietal lobule, may also be particularly susceptible to increases in arousal as noradrenergic projections from the LC are thought to underlie the TPJ’s attention-orienting function (Corbetta et al., 2008). Consistent with this theory, propranolol, a beta-adrenergic blocker blunts TPJ responses to novel stimuli (Strange & Dolan, 2007). Lesions of the TPJ have also led to reductions in the P300 event-related potential response, which is thought to reflect phasic activity of the noradrenergic system (Nieuwenhuis et al., 2005). However, it is not clear whether increases in emotional arousal would modulate TPJ responses associated with bottom-up attention to salient cues and whether it may contribute to age differences in arousal-induced selectivity.

Finally, we measured LC structure and function to test contributions of the LC to age differences in arousal-induced selectivity. To delineate the LC, a turbo-spin echo (TSE) T1-weighted MRI protocol was used, which visualizes the LC as a hyperintense region adjacent to the floor of the fourth ventricle (Sasaki et al. 2006). From these images, a semi-automatic procedure (Dahl et al. 2019) was used to compute contrast ratios as a measure of LC structure (Liu et al. 2017). To index LC function, coordinates from a preexisting atlas (Keren et al. 2009) were used to localize an LC ROI and pupil size was measured in response to the arousal manipulation (pupil dilation has been associated with LC BOLD activity; Murphy et al., 2014). As it is unknown whether LC structural MRI contrast covaries with LC function, we probed the relationship between these measures and their association with behavioral and fMRI outcomes.

## 2. Material and Methods

### 2.1 Participants

Participants were invited to participate if they were right-handed, had normal hearing, normal or corrected-to-normal vision, no history of psychiatric or neurological disorders, and were not taking beta-blocker medications. The final sample included 30 young adults aged 19-29 years (*M* = 23.43, *SD* = 2.83) and 30 older adults aged 60-85 years (*M* = 70.63, *SD* = 6.16). All participants received $75 USD for their participation. Three participants’ data were excluded and replaced: One young adult because of an incidental finding in their MRI images, one older adult because the MRI-compatible response pad malfunctioned, and one older adult whose data were not saved to the MRI scanner due to a technical issue.

Table 1 displays characteristics of the final sample. There were no age differences in sex ratio or education. On the 21-item Depression Anxiety Stress Scales (DASS-21; Lovibond & Lovibond, 1995), scores fell in the normal range with no significant age differences in depression or stress, but higher levels of anxiety in young than in older adults. Older adults also scored higher than young adults on the Shipley Vocabulary Test (Shipley, 1940). Participants were screened for cognitive impairment using the Montreal Cognitive Assessment (MoCA; Nasreddine et al., 2005). Scores were in the normal cognition range (>26) for both groups.

**Table 1.** Sample Characteristics

| Characteristic | Young Adults<br>M (SD) | Older Adults<br>M (SD) | Group Difference |
| --- | --- | --- | --- |
| Age | 23.43 (2.84) | 70.63 (6.16) | $t(58) = -38.14, p < .001$ |
| Sex ratio | 18F:12M | 15F:15M | $\chi^2(1) = .27, p = .60$ |
| Years of education | 16.77 (2.34) | 17.90 (3.10) | $t(58) = -1.59, p = .12$ |
| Vocabulary level <sup>a</sup> | 31.23 (3.41) | 35.50 (3.49) | $t(58) = -4.78, p < .001$ |
| Depression <sup>b</sup> | 6.07 (7.13) | 3.80 (5.64) | $t(58) = 1.36, p = .18$ |
| Anxiety <sup>b</sup> | 5.27 (5.19) | 2.80 (3.31) | $t(58) = 2.19, p = .03$ |
| Stress <sup>b</sup> | 9.53 (7.18) | 6.93 (5.22) | $t(58) = 1.61, p = .11$ |
| MoCA <sup>c</sup> | 28.50 (1.22) | 26.79 (2.23) | $t(56) = 3.66, p = .001$ |

*Note.* <sup>a</sup> Assessed with Shipley Vocabulary test (total possible = 40); <sup>b</sup> Assessed with 21-item Depression, Anxiety, and Stress Scales (DASS-21; total possible = 42 for each sub-scale); <sup>c</sup> Montreal Cognitive Assessment, normal cognition $\geq 26$ . F = female, M = male.

### 2.3 Materials

The experimental tasks were programmed in MATLAB using Psychophysics Toolbox Version 3. Stimuli included images of bodies and scenes to activate the EBA (Downing et al., 2001) and the PPA (Epstein & Kanwisher, 1998), respectively. For the incidental encoding task, 96 clothed body images (including 32 full body, 32 torso only, 32 legs only) and 96 scene images (48 indoor, 48 outdoor) were selected from previous studies (Downing et al., 2006; Kaiser et al., 2014; Orlov et al., 2010; Taylor et al., 2007), the SUN database (Xiao et al., 2010), and the Internet. Each image was yoked with an image from the same category that was perceptually different, which served as a foil during the recognition task. This resulted in a total of 192 images within each stimulus category. Images were resized to 300×300 pixels and normalized to the mean luminance of all images using the SHINE toolbox in MATLAB (Willenbockel et al., 2010). During the encoding task, stimuli sets were divided in two and counterbalanced as visually salient and non-salient images. Salience was manipulated on each trial by degrading the contrast of non-salient images by 80% and adding a yellow border to salient images. An additional novel 36 body images and 36 scene images were selected for a functional localizer task that was used to delineate the EBA and PPA.

Sound clips were used to induce arousal on a trial-by-trial basis. A set of 48 arousing and 48 non-arousing 6-second sound clips were selected from the International Affective Digitized Sounds database (Bradley & Lang, 2007). Using Audacity® recording and editing software (version 2.4.2; https://audacityteam.org/), sound clips were trimmed to include the most intense 3 seconds of each clip. Participants rated the 3-second sound clips using a 5-point Likert-style self-assessment manikin ranging from 1 = not arousing at all to 5 = very arousing.

### 2.3 Procedure

Participants provided informed consent prior to starting the experiment and procedures adhered to guidelines for ethical human subject research as approved by the Institutional Review Board at University of Southern California. At the beginning of the session, the DASS-21, Shipley Vocabulary Test, and a background questionnaire were administered using Qualtrics survey platform. The experimenter then conducted the MoCA. After this and before entering the scanner, 12 practice trials for the incidental encoding task were completed. The timing of each trial is depicted in Figure 1. Participants first heard a sound clip while they fixated on a cross at the center of the screen. A body and scene image then appeared side by side, one of which was salient and the other non-salient. A dot probe then replaced one of the images on either the left or right side of the screen and participants detected its location as quickly as possible by pressing a button with their index (left response) or middle finger (right response) on a fiber optic fMRI-compatible response pad with their right hand. Image selection was randomized, but each stimulus category was presented as salient or non-salient as well as on the left or right side of the screen equally across participants. The dot probe also appeared equally as often behind salient or non-salient images and on the left and right side of the screen.

**Figure 1.**
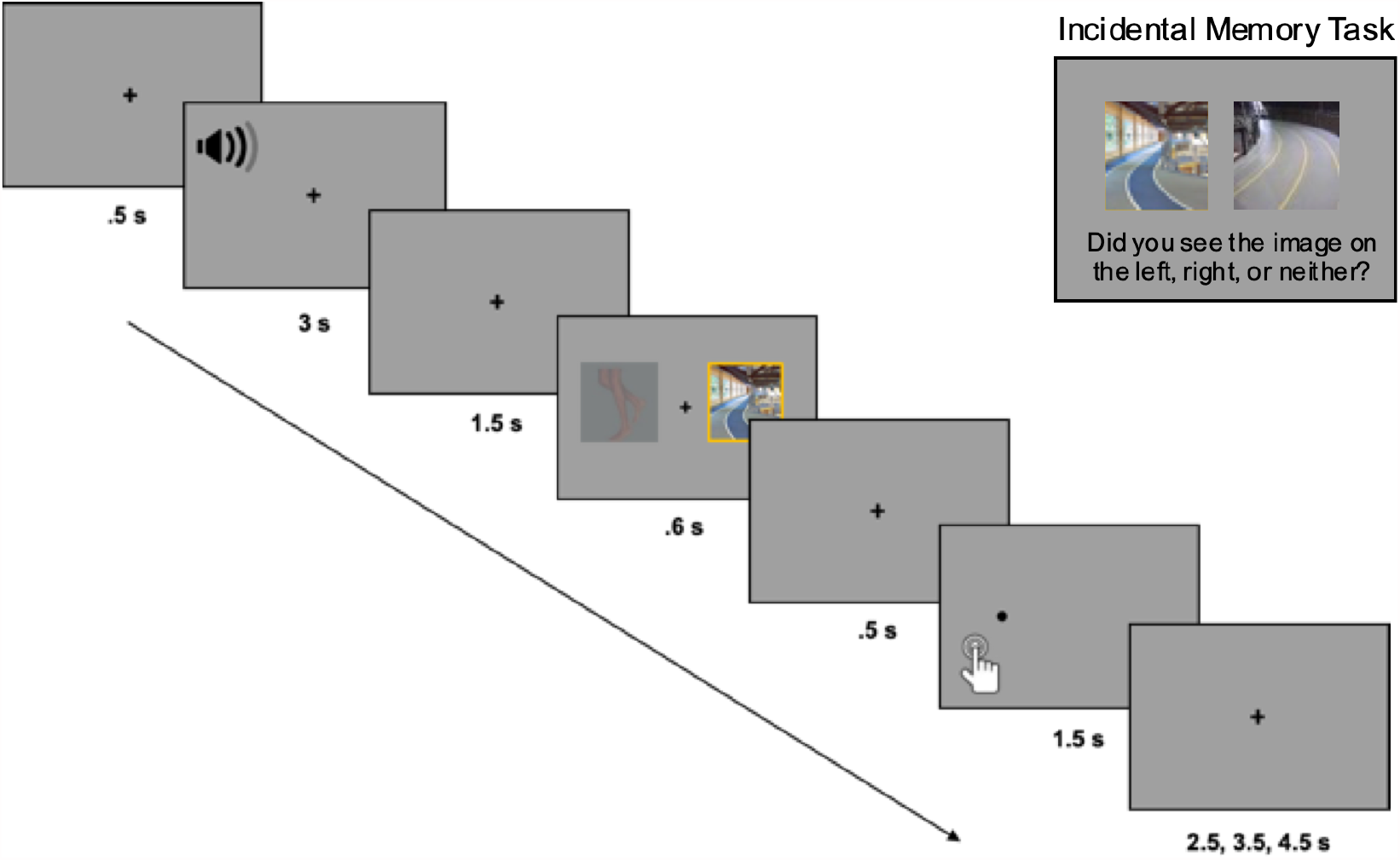
A sample trial from the dot-probe/encoding task and the incidental memory task.

The scanning protocol lasted approximately one hour. The high-resolution T1-weighted MPRAGE anatomical scan was acquired first followed by a T1-weighted TSE scan, which was used to delineate the LC in the brainstem. After structural scans, fMRI volumes were collected across three blocks of the encoding task (Figure 1), which each consisted of 32 trials. Sounds were played through Sensimetrics Model S14 MRI-compatible earbuds. Between each block, participants were reminded of the task instructions and to keep their head as still as possible. The entire task took ∼18 min to complete. The EBA/PPA functional localizer followed, which entailed a 1-back working memory task with a blocked design that included three body and three scene blocks of 14 trials each. Responses were made with the index finger of the right hand and were only required on 1-back trials of which there were two per block. Each trial started with a fixation cross for 500 ms followed by an image presented for 150 ms; blocks were separated by an 8-second fixation cross. To closely match the design of the encoding task, images appeared equally on the left or right side of the screen. The scanning session concluded with a 5-min resting state scan, the results of which are not included here.

Once outside of the scanner, participants completed a surprise recognition task. Memory was tested for all body and scene images presented during the encoding task. On each trial, a pair of images from the same stimulus category was presented and participants indicated whether they saw the image on the left, the image on the right, or neither of the images via keyboard press; responses were self-paced. There were 96 trials that contained an old image yoked with a lure image from the same stimulus category as well as 32 trials (16 bodies, 16 scenes) that contained a pair of lure images to garner an estimate of guessing (i.e., false alarms). Following the recognition task, the 96 sound clips were rated on arousal level. Pupil dilation was continuously measured during the rating task using a Gazepoint GP3 60 Hz research-grade portable eye tracker. Participants fixated on a blank screen while listening to a sound clip for 3 s after which they made a self-paced arousal judgment. Responses were made via mouse click using a 5-point self-assessment manikin ranging from 1 (not arousing at all) to 5 (very arousing).

### 2.4 MRI Data Acquisition

Neuroimaging data were acquired on a 3T Siemens Magnetom Prisma MRI system in the Center for Image Acquisition at USC Keck School of Medicine. Stimuli were displayed on a 32-inch Cambridge Research BOLDscreen 32 with a 1980×1080 pixel widescreen LCD display and a fixed 120 Hz frame rate, viewed via a mirror attached to a 32-channel matrix head coil. A high-resolution T1-weighted MPRAGE anatomical sequence was acquired first to help with functional image co-registration and LC delineation (TR = 2300 ms, TE = 2.26 ms, TI = 1060 ms, 176 interleaved slices, flip angle = 9°, FoV = 256mm, slice thickness = 1mm with no gap, bandwidth = 200 Hz/Px, GRAPPA acceleration factor = 2, 1mm isotropic voxel size) followed by a two-dimensional, multi-slice T1-weighted TSE sequence (TR = 750 ms, TE = 10 ms, 11 axial slices, flip angle = 120°, FoV = 220mm, bandwidth = 287 Hz/Px, slice thickness = 2.5mm, slice gap = 1.0 mm, in-plane resolution = 0.429mm^2^). Functional images for the localizer task (111 volumes) and three blocks of the encoding task (170 volumes each) were acquired using a multiband echoplanar sequence (TR = 2000 ms, TE = 54.60 ms, 88 interleaved slices with no gaps, flip angle = 52°, FoV = 222mm, bandwidth = 2228 Hz/Px, acceleration factor = 8, and 1.7mm isotropic voxel size).

### 2.5 Behavioral and Pupil Analyses

#### 2.5.1 Subjective arousal ratings and pupil dilation response to sound clips

We first examined the effect of sound type on subjective arousal ratings and pupil dilation during the rating task. To determine the effect of sound type on subjective arousal level, each participant’s ratings were averaged and analyzed in a 2 (age group: young, older) x 2 (arousal: arousing, non-arousing) mixed-effects ANOVA.

The pupillometry data were preprocessed using MATLAB (Mathworks, MA). Blinks and other artifacts were removed from the signal using a semi-automated program developed in our lab (github.com/EmotionCognitionLab/ET-remove-artifacts). The program detects blinks using an algorithm based on the velocity profile of each participants’ pupil time course (Mathôt et al., 2013) and linearly interpolates over the detected blink region. For several participants, the presence of non-blink artifacts impeded the precision of the blink-removal algorithm and so these time courses were manually corrected. Segments of non-blink artifacts that exceeded a duration of 1 s were classified as missing data and ignored in analyses.

For the pupil dilation analysis, we computed baseline pupil diameter and change in pupil diameter for each sound. If over half of the samples from a single trial were imputed with missing data indicators, then the trial was removed from analyses. After applying these exclusion criteria, the data from 25 older adults and 29 young adults were included in the pupil analysis. Average pupil dilation was examined in response to arousing and non-arousing sounds. For each trial, we searched for maximum pupil dilation in a window from the sound onset to 3 seconds after sound onset. To determine dilation over the trial, the peak pupil dilation in this time window was baseline normed by subtracting the average pupil dilation during the 500 ms before sound onset. Pupil dilation responses were then analyzed in a 2 (age group: young, older) x 2 (arousal: arousing, non-arousing) mixed-effects ANOVA.

We also examined correspondence between subjective and objective markers of arousal across age groups, specifically whether trial-by-trial pupil dilations predicted subjective ratings across arousal levels and age groups. Linear mixed-effect models (LMM) were used to model the data, implemented in R using the lme4 (Bates, Mächler, Bolker, & Walker, 2015) and lmerTest (Kuznetsova et al., 2017) packages. Subjective ratings were the dependent variable and arousal condition (coded as 1 = arousing, 0 = non-arousing), age group (coded as 1 = young, -1 = older), and continuous sound-evoked pupil dilation were modelled as fixed effects. Participant intercepts were modeled as random effects (i.e., random variance in intercepts due to unexplained differences between participants; Meteyard & Davies, 2020).

#### 2.5.2 Attentional bias and memory analyses

Attention bias and memory analyses were carried out separately for bodies and scene images to determine how arousal-induced selectivity differed as a function of distinct category-selective stimuli. To determine the effect of arousal on perceptual selectivity, we examined reaction times (RTs) during the incidental encoding task as a function of age group. The RTs from error trials as well as those ± 2.5 SDs from each participant’s average were removed prior to computing mean RTs for each condition. We computed attentional bias scores by subtracting the mean RT for probes appearing on the same side as a non-salient image from the mean RT for probes appearing on the same side as a salient image. Higher bias scores would indicate a bias to attend to salient images and lower values would indicate a bias to attend to non-salient images. We then compared attentional bias scores across age groups in a 2 (age group: older, young) x 2 (arousal: arousing, non-arousing) mixed-effects ANOVA and one-sample t-tests to determine whether age groups were more or less likely to attend to salient than non-salient images on arousing and non-arousing trials.

To determine age differences in arousal’s impact on memory selectivity, a 2 (age group: older, young) x 2 (arousal: arousing, non-arousing) x 2 (salience: salient, non-salient) mixed-effects ANOVA was conducted on mean hit rates to correctly identify old images as old. We also examined age differences in false alarm rates using an independent-samples t-test.

### 2.6 Neuroimaging Preprocessing and Analysis

Image preprocessing was carried out with FMRIB Software Library (FSL, www.fmrib.ox.ac.uk/fsl). Functional volumes were processed using FSL’s FMRI Expert Analysis Tool (FEAT, Version 6) with the following steps: motion correction using MCFLIRT, non-brain tissue removal using BET, spatial smoothing using a Gaussian kernel of 5mm full-width-at-half-maximum, grand-mean intensity normalization of the entire 4D dataset by a single multiplicative factor, and high-pass temporal filtering (Gaussian-weighted least-squares straight line fitting, with sigma = 50.0s). Independent component analysis (ICA) was carried out using MELODIC to identify unexpected artifacts or activation (Beckmann & Smith, 2004). To classify noise components, FMRIB’s ICA-based Xnoisefier (FIX) auto-classifier was run on single-subject ICA outputs. The FIX classifier was trained on 10 single-subject ICA datasets from the current study in which noise artifacts related to head motion, white matter/CSF signal, cardiac or respiratory pulsation, artery activation, or the multiband MRI-sequence were manually identified and removed (Griffanti et al., 2017). The auto-classifier then classified and removed noise components from the remaining 50 single-subject ICA datasets.

For whole-brain analysis, each participant’s denoised mean functional volume was registered to their T1-weighted anatomical image using boundary-based registration (BBR) with FSL’s FLIRT (Jenkinson et al., 2002). An affine transformation with 12 degrees of freedom was then used to register anatomical images to MNI-152 standard-space with 2mm resolution.

#### 2.6.1 Whole-brain voxel-wise analysis

A general linear model (GLM) was set up to examine processing of salient versus non-salient category-selective images on arousing compared to non-arousing trials. Event-related time vectors were created that modeled the onset times of the image pairs, with a duration of 600 ms, and were used as task regressors in the model. A lower-level GLM was built for each participant using four task regressors: (1) arousing salient body and non-salient scene, (2) arousing non-salient body and salient scene, (3) non-arousing salient body and non-salient scene, and (4) non-arousing non-salient body and salient scene. Each regressor was convolved with the canonical hemodynamic response function (HRF). Contrasts were created to test for main effects of arousal (arousing minus non-arousing), salience (salient minus non-salient), and arousal-by-salience interactions (arousing salient minus arousing non-salient, non-arousing salient minus non-arousing non-salient). A second lower-level GLM was conducted that modeled the onset times of the sound clips to better isolate the effects of arousal on LC function and its association with LC contrast and pupil dilation. Contrasts in this analysis tested for main effects of arousal (arousing minus neutral trials).

Second-level fixed-effects analyses averaged each participant’s functional runs. To determine group-level effects, the contrast images from the second-level analyses were analyzed in a higher-level random-effects analysis computed with fMRIB’s mixed-effects FLAME 1 model (Beckmann et al., 2003). Clusters were determined by thresholding the statistical images at Z > 2.3, corrected with a family-wise error rate (*p* < .05) at the cluster level.

#### 2.6.2 Regions of interest (ROI) analysis

Analyses were performed to determine age differences in how arousal modulated activity in the LC as well as the interaction of arousal and salience in category-selective visual cortex, and the TPJ. For the LC, we used a mask in 0.5mm MNI152 standard space from a previous study (Keren et al., 2009). To warp the mask to 2mm MNI152 standard space, we first co-registered the 0.5mm linear brain to MNI152 2mm standard space using the “antsRegistrationSyN.sh” routine in Advanced Normalization Tools (ANTS) v2.3.4 (Avants et al., 2009). The resulting transformations were then applied to warp the LC mask into 2mm space. Using this warped mask, percent signal change values were extracted from each participant’s functional scan using Featquery in FSL and submitted to a 2 (age group: young, older) x 2 (arousal: arousing, non-arousing) mixed-effects ANOVA.

Category-selective ROIs were functionally defined for each participant as a 6mm sphere centered on the peak voxel in the EBA and PPA that was most selective for bodies (bodies > scenes; *Z* = 2.3, uncorrected) and for scenes (scenes > bodies; *Z* = 2.3, uncorrected) during the localizer task. With this approach, we were able to define ROIs for all participants for the EBA (mean peak voxel in MNI: left [-48 -78 2]; right [54 -68 -4]) and PPA (mean peak voxel MNI coordinates: left [-22 -42 -12]; right [20 -34 -16]). The masks were then aligned to each participant’s second-level statistical parametric maps and used to extract percentage signal change values of encoding-related activity in the left/right EBA and PPA. To test our hypothesis that arousal would selectively enhance activation in body-selective and scene-selective voxels in young but not older adults, we performed separate 2 (age group: young, older) x 2 (arousal: arousing, non-arousing) x 2 (salience: salient, non-salient) x 2 (hemisphere: left, right) mixed-effects ANOVAs on the extracted means from each ROI.

Next, we examined age differences in the arousal-by-salience interaction in the right TPJ, a node of the ventral attention network involved in salience-driven attention to stimuli (Kahnt & Tobler, 2013). We focused on the right TPJ given evidence that the ventral attention network is largely lateralized to the right hemisphere (Fox et al., 2006). The TPJ mask was created using a connectivity-based parcellation atlas (Mars et al., 2012), which includes anterior and posterior TPJ subdivisions in the right hemisphere. These clusters were combined and binarized using fslmaths in FSL. The final mask extended from MNI coordinates [68 -31 22] to [60 -61 22]. Percent signal change values of encoding-related activity were then extracted from the right TPJ and analyzed in a 2 (age group: young, older) x 2 (arousal: arousing, non-arousing) x 2 (salience: salient, non-salient) mixed-effects ANOVA.

#### 2.6.3 Calculation of LC contrast ratios

An automated approach was used to delineate the LC on TSE scans. This entailed bringing participants’ TSE scans into MNI-ICBM 152 0.5 mm linear space using routines within ANTS v. 2.3.4 (Avants et al., 2009). A previously validated approach was used (Dahl et al., 2019), detailed in the Supplementary Methods.

Using the TSE template generated during LC delineation (warped to MNI 0.5mm linear space), we confirmed that at a group level, TSE hyperintensities fell within the LC map (Keren et al., 2009) used for functional analyses (Figure 2). We applied this LC mask on individual TSE scans in MNI 0.5mm linear space to isolate probable LC voxels for contrast ratio calculation. In line with the anatomical boundaries of the LC, we restricted the search for probable LC voxels to MNI z = 85 to 112 (Ye et al., 2021). For each participant, the intensity of the peak voxel across all z-slices within the masked region was extracted for each hemisphere separately. We then masked each participant’s TSE scan in MNI 0.5mm linear space with a publicly available 4×4mm reference region covering the central pontine region (Dahl et al., 2020) and extracted the peak intensity value across all z-slices in the masked region. LC contrast ratios were calculated from peak intensities as follows (Liu et al., 2017): 

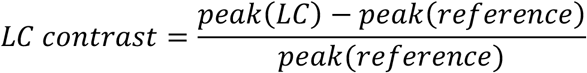

**Figure 2.**
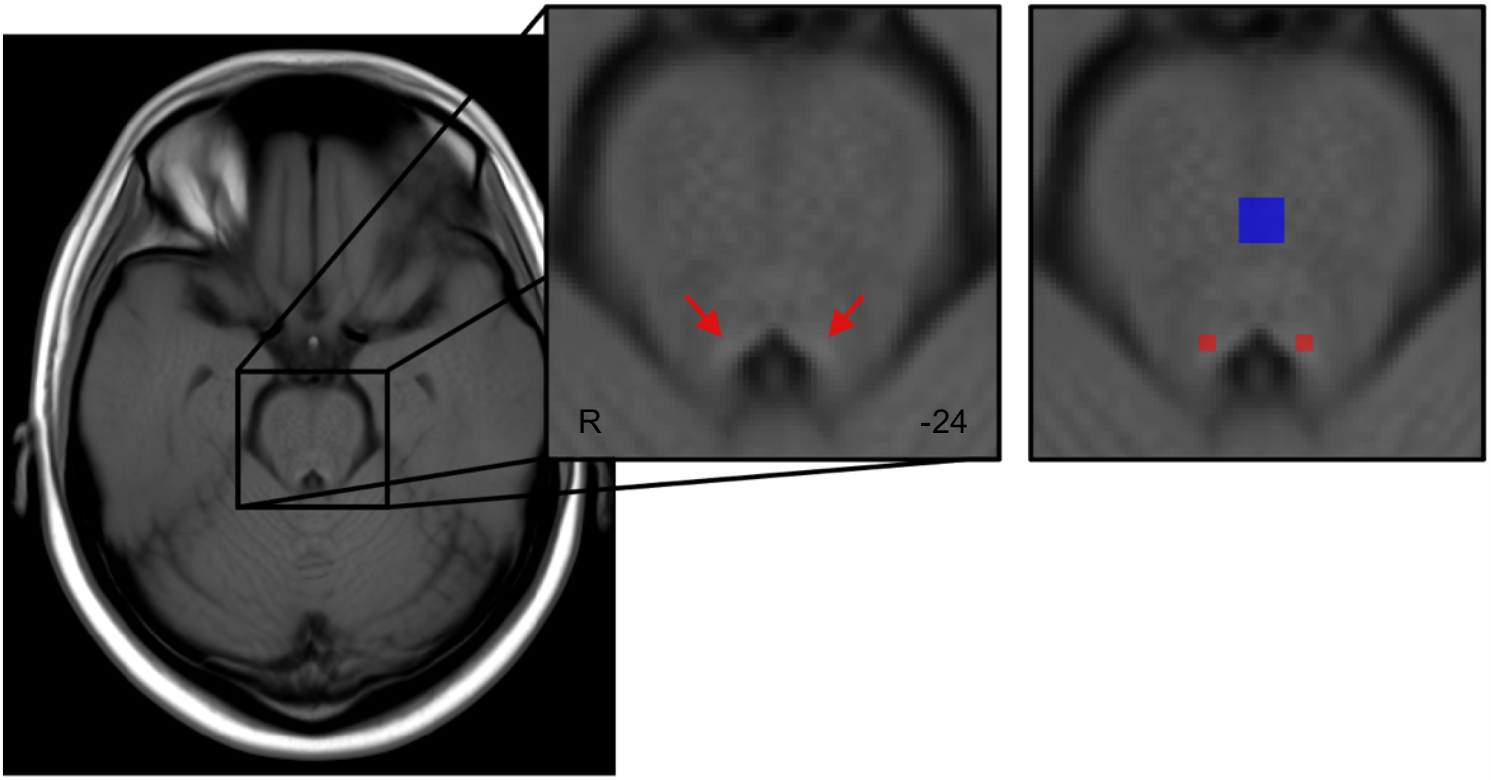
Template of all participants’ TSE scans, warped to MNI152 0.5mm linear space in the axial plane. Red arrows indicate hyperintensities bordering the fourth ventricle. For calculation of LC contrast ratios, a previously published LC map (Keren et al., 2009) was applied as a mask on individual TSE scans (shown in red) to identify probable voxels corresponding to the LC. In addition, a reference region covering the central pons (Dahl et al., 2020) was applied as a mask on individual TSE scans (shown in blue). Peak intensities within the masked LC and reference regions were extracted for each participant to calculate contrast ratios.

Resulting LC ratios were analyzed in a 2 (age group: young, older) x 2 (hemisphere: left, right) mixed-effects ANOVA. One participant was excluded from this analysis because a susceptibility artifact was overlapping the pons in their TSE scan. To validate this automated approach, we used a manual protocol for assessing LC intensity (see Liu et al., 2017). We calculated the consistency of resulting LC peak intensities across methods via two-way mixed intraclass correlation coefficients (ICC), which are reported in the Supplementary Results.

#### 2.6.4 Associations between LC function, LC contrast, and pupil dilation

Finally, we used Pearson’s correlation analyses to probe relationships between LC contrast, LC function, pupil dilation, and behavior in each age group. To examine relationships with behavior, a selectivity index was computed as recognition of salient images minus recognition of non-salient images separately for arousing and non-arousing trials. Fisher’s *r*-to*-z* transformations were used to compare correlation coefficients between age groups.

## 3. Results

### 3.1 Behavioral and Pupillometry Results

#### 3.1.2 Subjective arousal ratings and pupillary responses to sounds

Subjective arousal ratings were higher for arousing (*M* = 3.82, *SD* = .95) than non-arousing sounds (*M* = .73, *SD =* .24), *F*(1, 57) = 572.86, *p* < .001, η_p_^2^ = .91, and older adults showed overall higher ratings than young adults (*M*_*older*_ = 2.42, *SD* = .33 vs. *M*_*young*_ = 2.14, *SD* = .52), *F*(1, 57) = 5.72, *p* = .02, η_p_^2^ = .09. A significant interaction between age group and arousal, *F*(1, 57) = 7.04, *p* = .01, η_p_^2^ = .11, on subjective ratings further showed arousing sounds were rated as higher in arousal by older adults (*M* = 4.14, *SD* = .75) than by young adults (*M* = 3.52, *SD* = 1.04), *p* = .01, 95% CI [-1.089, -.144]. There was no age difference in ratings for non-arousing sounds (young: *M* = .77, *SD* = .23, older *M* = .69, *SD* = .25), *p* = .26, 95% CI [-.054, .195]. Similarly, pupil responses to were greater for arousing than non-arousing sounds, *F*(1, 52) = 56.36, *p* < .001, η_p_^2^ = .52, but there were no other significant effects or interactions on pupil responses, *Fs* ≤ 2.55, *ps* ≥ .12.

The LMM analyses queried the relationship between age group, pupil dilation, and sound type on arousal ratings (Table 2). Mirroring the ANOVA results on ratings, there was a main effect of arousal as well as a significant interaction of age group and arousal showing that older adults’ ratings were higher than young adults for arousing sounds (Figure 3A). There was also an interaction of arousal and pupil dilation on ratings. Plotting this interaction showed that pupil dilation to arousing sounds were more positively associated, *β* = 137.75, 95% *CI* [13.07, 262.44], *t* = 2.17, *p* = .03, with subjective ratings than pupil dilation to non-arousing sounds, *β* = 3.09, 95% *CI* [-78.44, 140.22], *t* = 0.55, *p* = .58 (Figure 3B).

**Table 2.** Linear Mixed-Effects Models Probing the Relationship Between Age Group, Arousal Condition, and Pupil Dilation on Subjective Arousal Ratings for Sound Clips

| Predictors | Estimate | S.E. | Wald's 95% CI | t-value | p-value |
| --- | --- | --- | --- | --- | --- |
| Intercept | 1.66 | 0.08 | 1.50, 1.82 | 20.46 | < .001 |
| Age | -0.06 | 0.08 | -0.22, 0.10 | -0.70 | 0.484 |
| Arousal | 2.16 | 0.029 | 2.10, 2.22 | 72.95 | < .001 |
| Pupil dilation | -20.12 | 64.24 | -145.98, 105.74 | -0.31 | 0.754 |
| Age x Arousal | -0.28 | 0.03 | -0.34, -0.23 | -9.59 | < .001 |
| Age x Pupil dilation | 1.95 | 64.24 | -123.91, 127.81 | 0.03 | 0.976 |
| Arousal x Pupil dilation | 294.39 | 90.28 | 117.45, 471.35 | 3.26 | 0.001 |
| Age x Arousal x Pupil dilation | 25.36 | 90.28 | -151.59, 202.34 | 0.28 | 0.779 |

*Note.* 95% confidence intervals were computed using `confint.merMod` function from the `lme4` library in R. Age (1=young, -1=older), arousal (1=arousing, 0=non-arousing), and pupil dilation to sounds were modeled as fixed effects. The participant intercept was modeled as a random effect.

**Figure 3.**
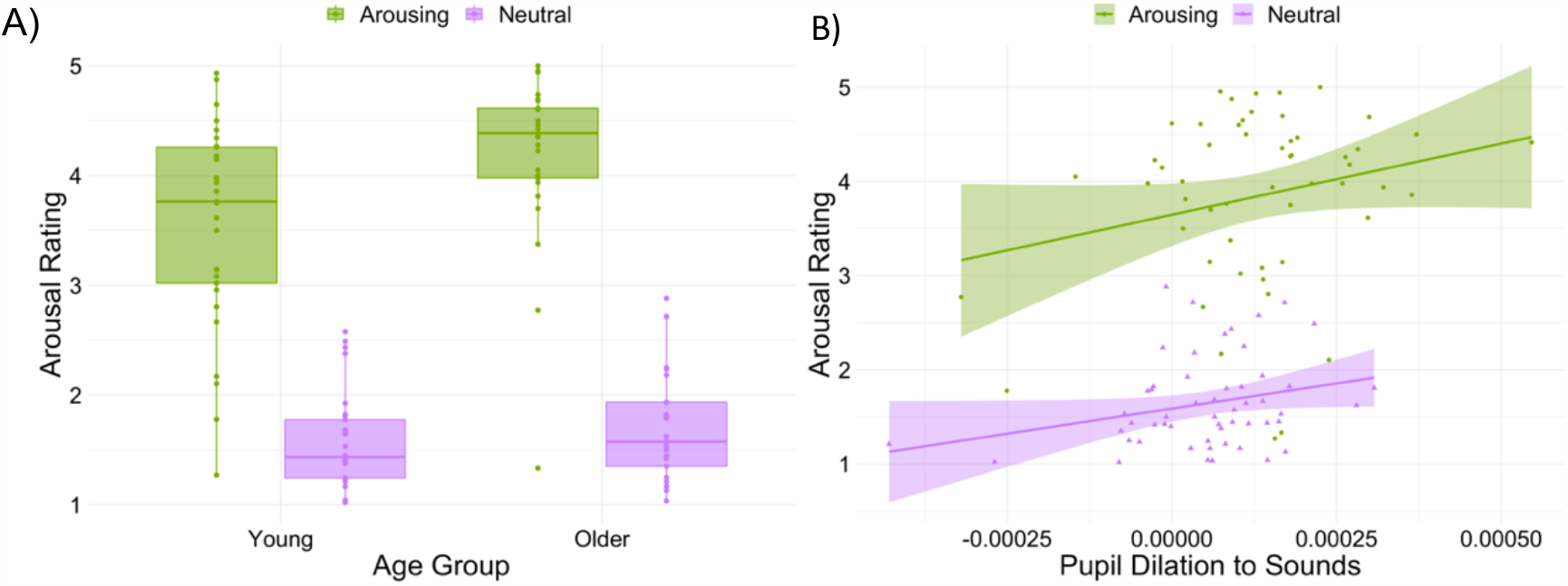
Results from linear mixed-effects model of subjective and objective arousal responses to sounds. A) Boxplots display the age x arousal interaction. Crossbars represent medians and the upper and lower hinges correspond to first and third quartiles, respectively. B) A significant arousal x pupil interaction showed that pupil dilation evoked by arousing sounds were more associated with subjective ratings than pupil dilation evoked by non-arousing sounds. Lines and shaded areas represent regression fits and standard error, respectively. For both plots, each dot represents a participant.

#### 3.1.3 Dot-probe attention bias scores

^1^ Mean reaction times (RTs) are presented in Table 3. These RTs were used to calculate attention bias scores for each arousal and stimulus condition (RT on non-salient probe trials minus RT on salient probe trials). The ANOVA on bias scores for body images showed an age-by-arousal interaction, *F*(1, 59) = 5.75, *p* = .020, η_p_^2^ = .09, suggesting that arousal had a different effect on bias scores in young and older adults. According to one-sample t-tests on their bias scores, young adults were slow to detect a salient probe on arousing trials, *t*(29) = -2.60, *p* = .014, *d =* -0.48, whereas a trending non-significant *t*-test showed that older adults were slow to detect salient probes on non-arousing trials, *t*(29) = - 1.89, *p* = .06, *d* = -0.34. There were no main effects or interactions in the analysis of attention bias scores for scene images, *Fs* ≤ 3.09, *ps* ≥ .09.

**Table 3.** Average Reaction Times (RT) for Detecting Probes

|  | Young |  | Older |  |
| --- | --- | --- | --- | --- |
|  | Arousing<br>M (SD) | Non-arousing<br>M (SD) | Arousing<br>M (SD) | Non-arousing<br>M (SD) |
| Salient Body RT | .398 (.09) | .395 (.09) | .554 (.13) | .575 (.13) |
| Non-salient Body RT | .380 (.09) | .390 (.10) | .556 (.13) | .559 (.13) |
| Salient Scene RT | .386 (.08) | .397 (.09) | .561 (.14) | .571 (.15) |
| Non-salient Scene RT | .385 (.10) | .383 (.08) | .549 (.13) | .572 (.14) |

#### 3.1.4 Incidental memory performance

^**2**^ Participants remembered more body images from arousing than non-arousing trials, *F*(1, 58) = 4.35, *p* = .04, η_p_^2^ = .07, and remembered marginally more salient than non-salient body images, *F*(1, 58) = 3.71, *p* = .06, η_p_^2^ = .06. Arousal had an opposing effect on memory selectivity across age groups as evidenced by an interaction of age, arousal, and salience, *F*(1, 58) = 5.52, *p* = .02, η_p_^2^ = .09 (Figure 4). Bonferroni-corrected pairwise comparisons showed that, for young adults, hit rates were higher for salient than for non-salient body images on arousing trials, *p* = .04, 95% *CI* [.003, .102], but not on non-arousing trials, *p* = .52, 95% *CI* [-.027, .053]. By contrast, for older adults, hit rates were higher for salient than for non-salient body images on non-arousing trials, *p* = .04, 95% *CI* [.002, .083], but not on arousing trials, *p* = .53, 95% *CI* [-.065, .034]. No other main effects or interactions were significant, *Fs* ≤ 3.21, *ps* ≥ .08. There was also no age difference in false alarm rates for body images (*M*_*young*_ = .39, *SD* = .29; *M*_*older*_ = .34, *SD* = .22) *t*(58) = .92, *p* = .36, *d* = .19.

**Figure 4.**
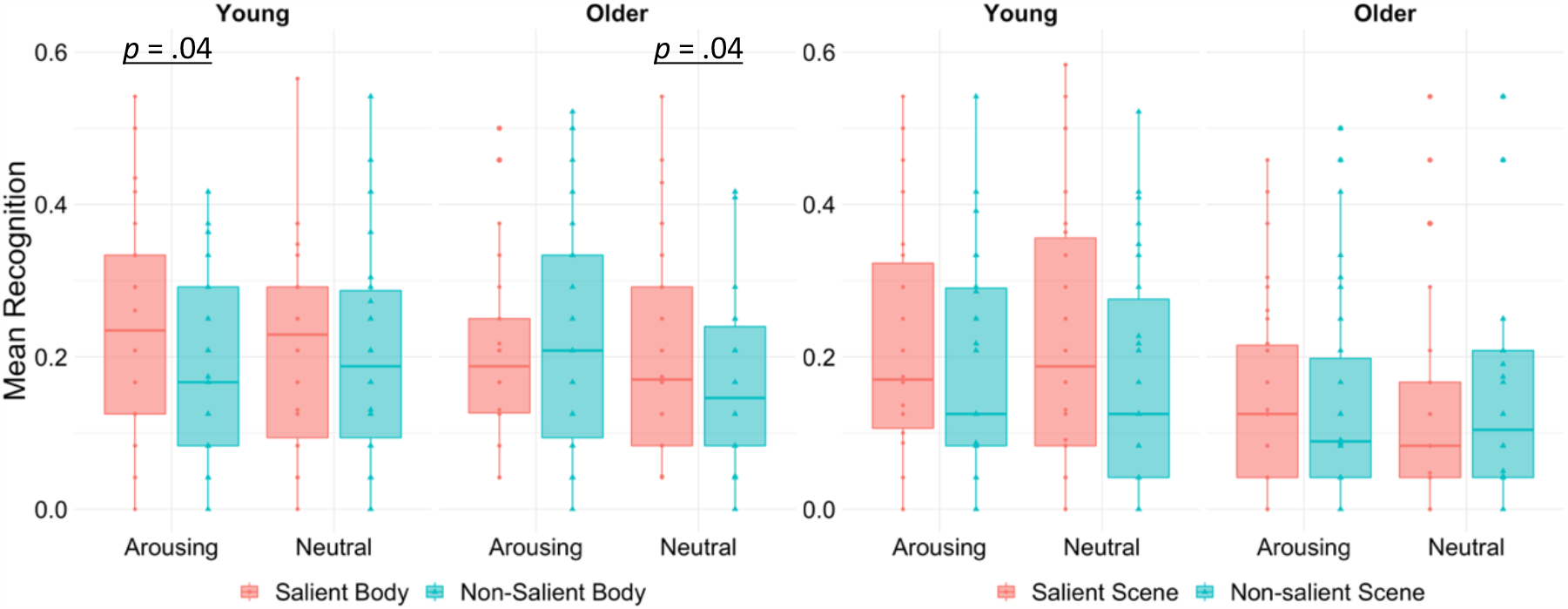
Box plots display memory performance for body and scene images as a function of age group, arousal, and salience. Crossbars represent medians and the upper and lower hinges correspond to first and third quartiles, respectively. Each dot represents a single participant.

For scenes, a trending non-significant effect of salience, *F*(1, 58) = 3.50, *p* = .06, η_p_^2^ = .06, showed that hit rates were numerically higher for salient than non-salient images (Figure 4). No other effects were significant, *Fs* ≤ 2.93, *ps* ≥ .09, nor was there an age effect on false alarms for scenes (*M*_*young*_ = .33, *SD* = .29; *M*_*older*_ = .21, *SD* = .23) *t*(58) = 1.77, *p* = .08, *d* = .47.

### 3.2 Brain Activity Results

#### 3.2.1 Whole-brain activation during the encoding task

Analysis of the BOLD signal at the whole-brain level revealed clusters showing a significant main effect of arousal on brain activity when participants were viewing the image pair. Arousing sounds increased activity in the frontal orbital cortex, supramarginal gyrus, superior and middle temporal gyrus, superior frontal gyrus, occipital fusiform gyrus, and the posterior cingulate gyrus. There was also an age-by-arousal interaction on encoding-related activity such that young and older adults showed different patterns of brain activity on arousing relative to non-arousing trials. In young adults, arousing trials corresponded with increased activation of frontal and temporal regions including the superior frontal gyrus, temporal pole, and the middle temporal gyrus, whereas older adults showed greater activity in a more posterior region of the superior frontal gyrus as well as in the superior parietal lobule, middle frontal gyrus, and precuneus cortex (Figure 5 and Table 4).

**Table 4.**
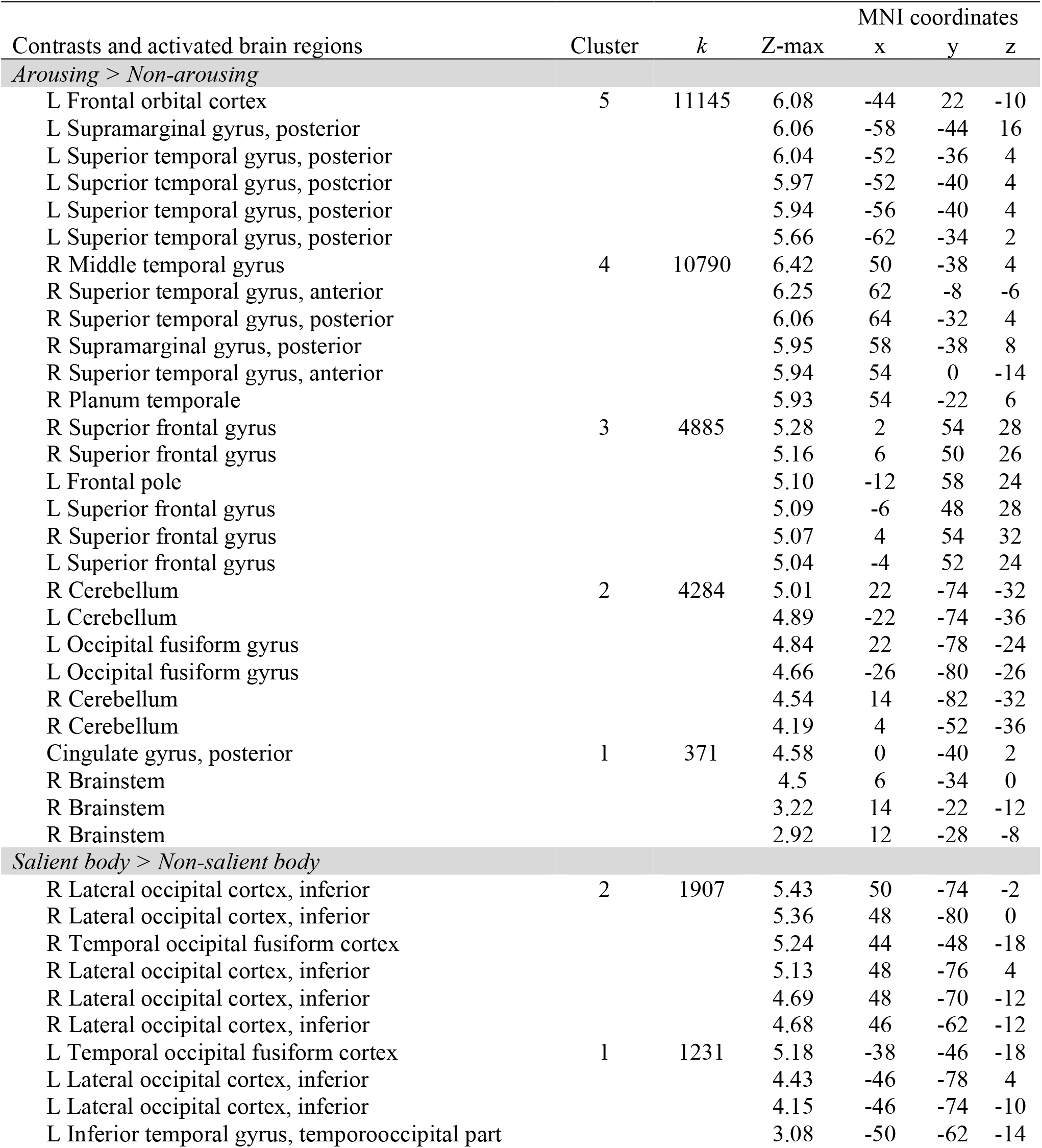
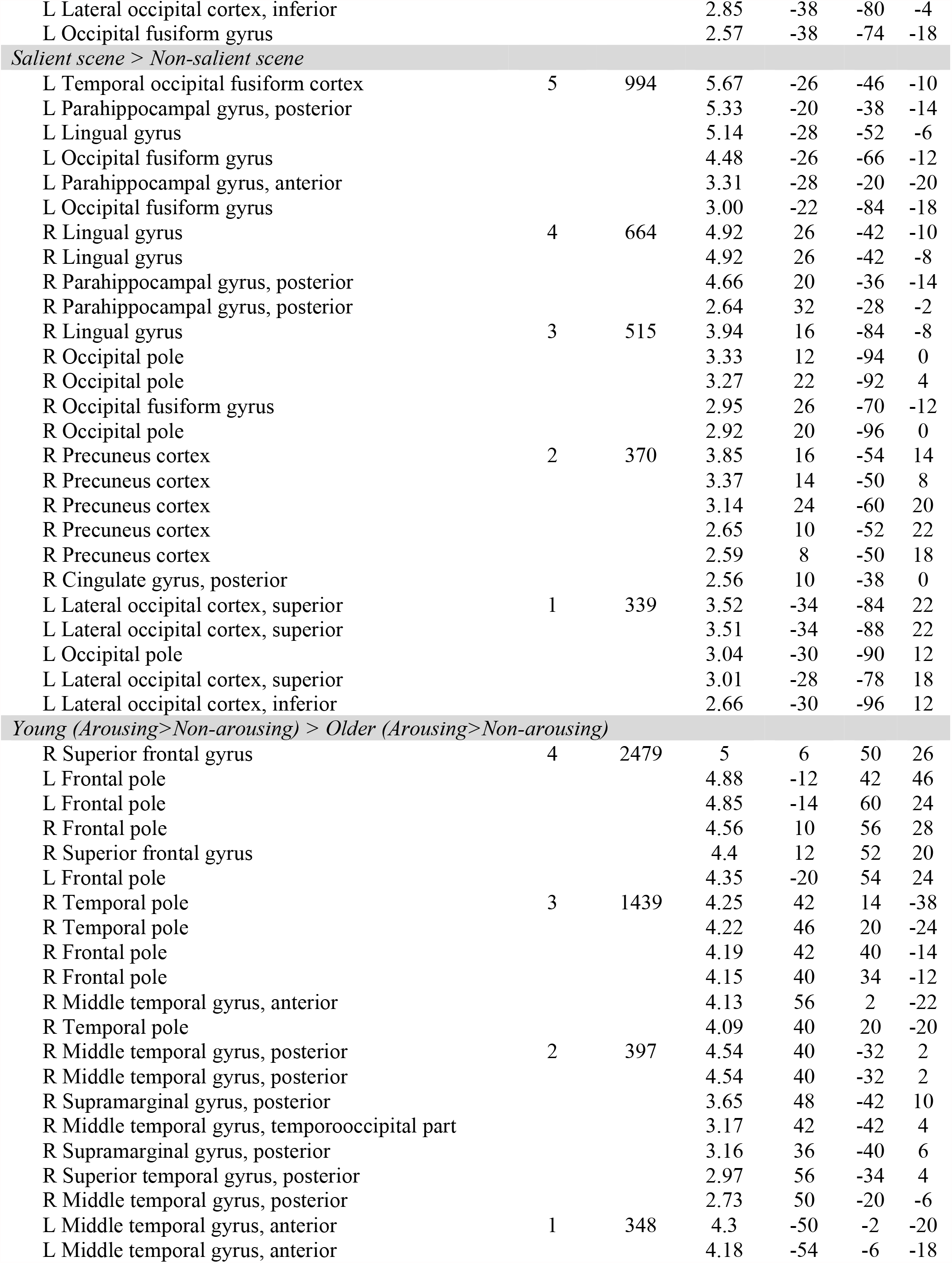
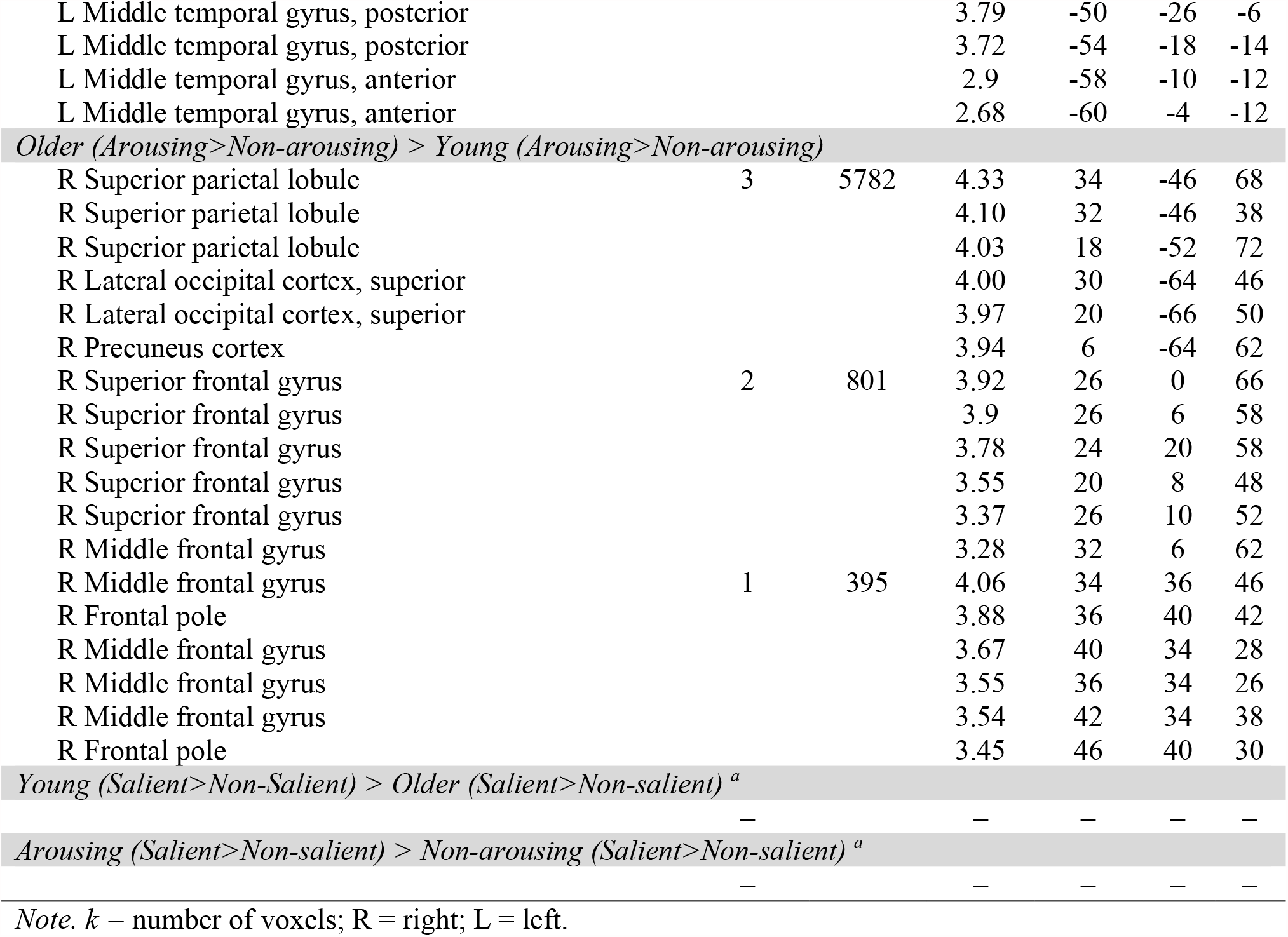
Significant Clusters and Locations of Local Maxima from the Whole-Brain Analysis of BOLD Signal when Viewing Images During the Encoding Task

**Figure 5.**
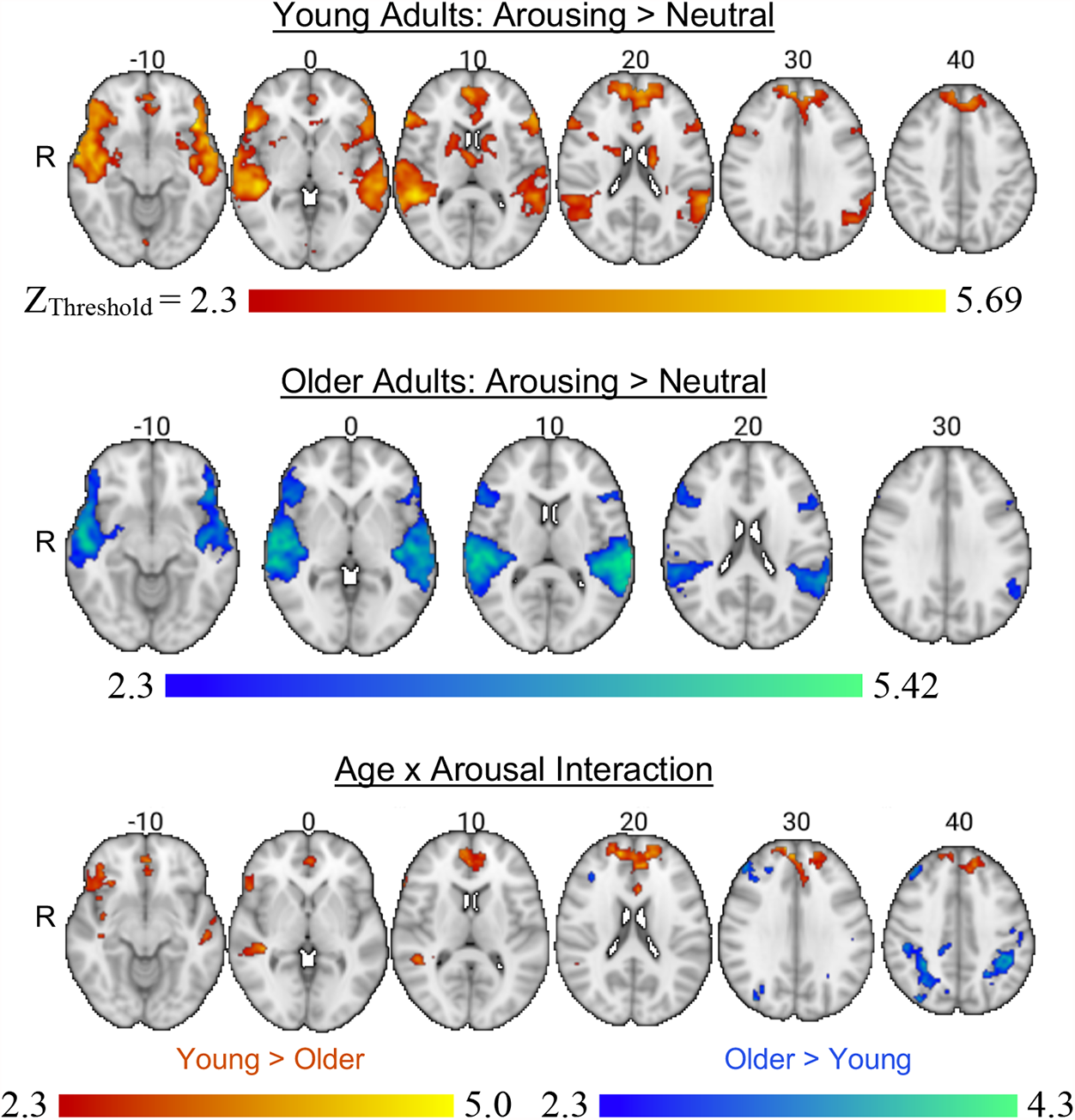
Whole-brain analysis results illustrating the effects of arousing sounds on brain activity when viewing images during the encoding task. The age x arousal interaction displays brain regions where age groups differed in the contrast of arousing versus non-arousing trials.

There were main effects of salience for both body and scene images, confirming that the manipulation of perceptual contrast worked. When participants were viewing salient relative to non-salient body images, brain activity was greater in occipitotemporal regions consistent with the location of the EBA (Downing et al., 2001) including the right and left lateral occipital cortex, temporal occipital fusiform cortex, and inferior temporal gyrus. Similarly, when contrasting salient scene images with non-salient scene images, significant clusters emerged in the parahippocampal gyrus, the temporal occipital fusiform cortex, lingual gyrus, occipital pole, and precuneous cortex. These effects of salience did not differ as a function of age group or arousal level as no significant clusters were observed for these contrasts (Table 4).

#### 3.2.2 ROI analysis results

Next, we performed analyses to determine how arousal modulated activity in our ROIs including the LC, category-selective occipitotemporal regions, and the right TPJ. For the LC ROI, there was a main effect of arousal, *F*(1, 58) = 4.17, *p* = .046, η_p_^2^ = .07, with higher activity on arousing than non-arousing trials. There was no significant effect of age group nor an interaction between age group and arousal, *Fs* ≤ .24, *ps* ≥ .63.

According to a main effect of salience, *F*(1, 58) = 33.62, *p* < .001, η_p_^2^ = .37, EBA activity was enhanced for salient body images relative to non-salient body images. An interaction of age, arousal, and salience, *F*(1, 58) = 7.79, *p* = .007, η_p_^2^ = .12, showed that the effect of arousal on EBA activation toward salient versus non-salient body images differed between age groups (Figure 6). As expected, Bonferroni-corrected pairwise comparisons revealed that EBA activation in young adults was greater for salient than non-salient body images on arousing trials, *p* < .001, 95% *CI* [.041, .124], but not on non-arousing trials, *p* = .36, 95% *CI* [-.029, .077]. Older adults’ brain activity showed the opposite pattern: EBA activity was greater for salient versus non-salient body images on non-arousing trials, *p* < .001, 95% *CI* [.063, .169], but not on arousing trials, *p* = .094, 95% *CI* [-.006, .076]. These effects did not differ by hemisphere, *F* = 1.09, *p* = .30. Similar to body images, salient scene images evoked more PPA activity than non-salient images, *F*(1, 58) = 29.92, *p* < .001, η_p_^2^ = .34. There was also a main effect of hemisphere, *F*(1, 58) = 11.52, *p* = .001, η_p_^2^ = .16, with greater activation in the right versus left PPA, but there were no other main effects or interactions, *ps* ≥ .05.

**Figure 6.**
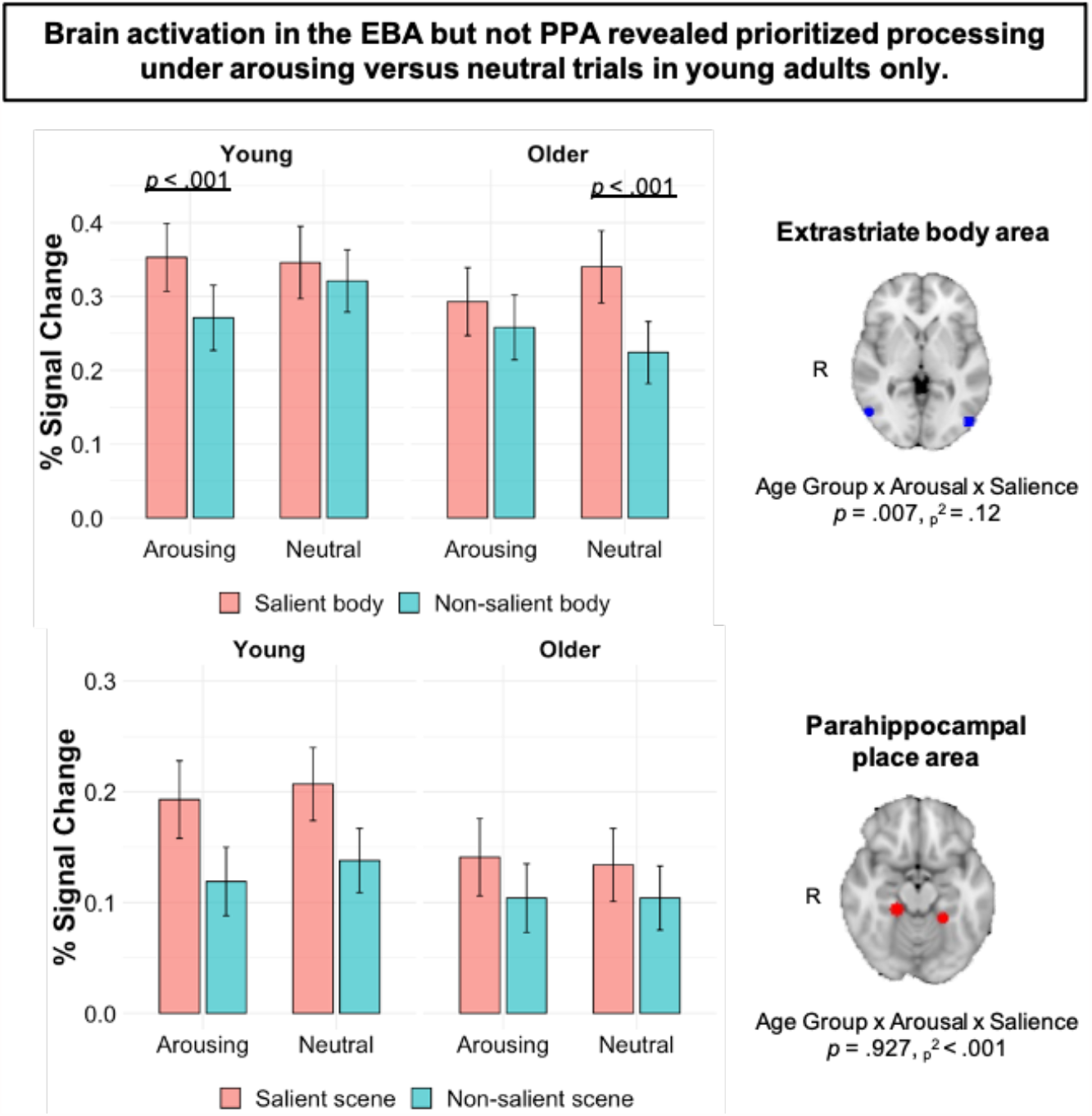
Mean percent signal change ± SEM in the EBA and PPA while viewing image pairs during the encoding task. The significant age x arousal x salience interaction for the EBA shows that arousing sounds enhanced prioritized processing in young adults, whereas older adults showed prioritized processing on trials with non-arousing sounds. There was no evidence for arousal-modulated processing in the PPA.

Finally, we examined age differences in the interaction of arousal and salience on activity in the right TPJ (rTPJ) in young and older adults. Given that we did not observe a three-way interaction in the behavioral and brain results for scene images, we focused this analysis on activity in the rTPJ when participants were encoding salient relative to non-salient body images. The analysis revealed a main effect of arousal, *F*(1, 58) = 23.46, *p* < .001, η_p_^2^ = .29, with significantly higher activity in the rTPJ on arousing trials versus non-arousing trials. There was also a significant three-way interaction between age, arousal, and salience, *F*(1, 58) = 4.93, *p* = .03, η_p_^2^ = .08. For young adults, pairwise comparisons showed increased activity in the rTPJ when encoding salient body images on arousing versus non-arousing trials, *p* = .021, 95% *CI* [.047, .133]. For older adults, the effect was reversed: Activity in the rTPJ was enhanced for non-salient body images on arousing versus non-arousing trials, *p* = .007, 95% *CI* [.016, .094]. In other words, arousal amplified activity in the rTPJ when young adults were encoding salient versus non-salient body images, whereas no such modulation of rTPJ activity by arousal was observed for older adults. There were no other significant main effects or interactions, *ps* ≥ .17.

#### 3.2.3 LC contrast ratios

The LC contrast ratios did not differ between young (*M* = .08, *SD* = .04) and older adults (*M* = .08, *SD* = .05), *F* = .01, *p* = .92, but the ratios were higher in the left hemisphere (*M* = .10, *SD* = .05) than in the right hemisphere (*M* = .06, *SD* = .04), *F*(1, 57) = 41.74, *p* < .001, η_p_^2^ = .42. This replicates a lateralized difference (left > right) in LC ratios that has been previously reported (Betts et al., 2017; Dahl et al., 2019).

#### 3.2.4 Relationship between LC structure, LC function, pupil dilation, and behavior

Pearson’s correlation analysis showed that LC contrast ratios, averaged across hemispheres, were negatively associated with LC function on arousing trials in older adults, *r*(29) = -0.415, *p* = .025, but not in young adults, *r*(30) = -0.066, *p* = .728. According to Fisher’s *r*-to*-z* transformation, the age difference between these correlation coefficients was not statistically significant, *Z* = -1.37, *p* = .085. The association between LC contrast ratios and LC function (both averaged across hemispheres) on neutral trials was not statistically significant in older adults, *r*(29) = -0.317, *p* =.094, or in young adults, *r*(30) = 0.116, *p* = .542.

For both age groups, associations between LC contrast ratios, averaged across hemispheres, and pupil dilation were not statistically significant on arousing trials (older adults: *r*(23) = -0.18, *p =* .42; young adults: *r*(29) = 0.06, *p* = .76) or on neutral trials. LC function was positively associated with pupil dilation on arousing trials in young adults, *r*(29) = 0.51, *p* = .005, but not in older adults, *r*(23) *=* 0.20, *p* = .36. Fisher’s *r*-to*-z* transformation indicated that these correlation coefficients were not statistically different across age groups, *Z* = -1.24, *p* = .11. There was no association between LC function and pupil dilation on neutral trials (older adults: *r*(23) = 0.18, *p =* .41; young adults: *r*(29) = 0.16, *p* = .41).

Finally, we tested the relationship between LC measures and memory selectivity. LC contrast ratios, averaged across hemispheres, were not associated with memory selectivity for body images on arousing trials (older adults: *r*(28) = 0.08, *p =* .67; young adults: *r*(30) = 0.13, *p* = .48), but they were positively associated with memory selectivity for body images on neutral trials in young, *r*(30) = .41, *p* = .03, and not older adults, *r*(28) = 0.19, *p* = .33. Fisher’s *r*-to*-z* transformation indicated that these correlation coefficients were not statistically different across age groups, *Z* = 0.88, *p* = .19. There were no statistically significant correlations between LC contrast and memory selectivity for scene images in either age group, *ps* > .38. There were also no statistically significant correlations between LC function and memory selectivity, *ps* > .21.

## 4. Discussion

In challenging or threatening situations, survival can depend on our ability to orient attention toward salient or advantageous stimuli. Previous research has shown that the way attention is modulated by the arousal system changes with age (Lee et al., 2018). This is thought to be driven by a combination of age-related changes in critical attention networks in the brain (Campbell et al., 2012; Kennedy & Mather, 2019) as well as in the noradrenergic system, including loss of LC integrity (Robertson, 2013) and increases in certain noradrenergic neurotransmitters (Mather, 2020). In the current fMRI study, we investigated age differences in the interaction of arousal and bottom-up (stimulus-driven) attention, incidental memory, and brain activity. Our results revealed age differences in how arousal modulates bottom-up attention to salient stimuli. In line with our hypotheses, arousal increased selective processing in young but not older adults. Interestingly, these selectivity effects were evident for body and not scene images, which may be related to the biological relevance of body images.

Perceptual selectivity was indexed by attentional bias scores based on RTs to detect probes appearing behind salient versus non-salient images. On arousing trials, young adults’ attention bias scores indicated that they were *slower* to detect probes appearing behind salient versus non-salient body images than they were on non-arousing trials. Why did arousal not facilitate attention by leading to quicker probe detection behind salient images? When a salient stimulus wins the competition for attention, it should lead to a favoring of stimuli appearing in the same location. Theoretically, an increase in arousal should enhance this bias (Mather & Sutherland, 2011), but the facilitation of responses at a salient location only lasts for a short time window of ∼300 ms. After this, responses typically become slower at the salient location, reflecting an inhibition of return (IOR) effect (Posner & Cohen, 1984). An IOR effect occurs when attention is initially oriented towards a salient or cued location but then is disengaged after a passage of time. The result is an inhibitory aftereffect where responding is delayed to information subsequently presented at the cued location (Klein, 2000). In the current study, there was a 500 ms interstimulus interval between the images and dot probe, suggesting an IOR effect may have occurred. It is possible that young adults’ attention was initially more captured by salient than non-salient body images and then disengaged during the 500 ms ISI, leading to slower probe detection in the location of the salient image. Our results also suggest that this potential IOR effect was enhanced on arousing trials. Further research, including a shorter ISI comparison condition, is needed to test this hypothesis that arousal modulates the IOR effect.

In terms of memory performance, we found that arousing trials amplified incidental memory for salient versus non-salient body images in young but not older adults. Hearing an arousing sound was not advantageous for older adults’ memory selectivity; instead, they showed better memory for salient than non-salient body images after hearing a non-arousing sound. This age difference is consistent with evidence that arousal is less effective at modulating processing of high over low priority information during aging (Lee et al., 2018). Thus, relative to young adults, arousing conditions may increase older adults’ tendency to process all stimuli indiscriminately of its relevance. This, in turn, could make older adults less able to suppress attention to distracting or goal-irrelevant information when experiencing increases in arousal (Durbin et al., 2018; Gallant et al., 2020). Our results add to these findings by showing that, relative to young adults, emotional arousal is less likely to improve older adults’ ability to attend to and remember the salient features of stimuli.

Results from the ROI analyses aligned with the behavioral findings. Analysis of category-selective cortex allowed us to compare how arousal changes the neural representation of salient information in an isolated brain region in young and older adults. When young adults were viewing perceptually salient versus non-salient body images, arousing sounds increased activity in the extrastriate body area, a region of the occipitotemporal cortex that responds selectively to images of human bodies (Downing et al., 2001). Older adults, instead, showed enhanced EBA activity for salient versus non-salient body images after hearing a non-arousing sound. A similar age difference in arousal-induced selectivity was observed in the rTPJ, a node of the ventral attention network. Relative to non-arousing trials, young adults’ rTPJ activity was enhanced for salient body images on arousing trials, whereas older adults’ rTPJ activity was enhanced for non-salient body images. These findings support the GANE model (Mather et al., 2016) by showing that increases in emotional arousal can amplify selective neural processing in young adults. Moreover, the results highlight how arousal-induced selectivity extends to instances where priority is determined by exogenous factors like perceptual salience, leading to selective modulation of ventral attention network regions. For older adults, however, arousal does not seem to have an advantageous effect in terms of their behavioral or neural selectivity.

Prior studies have shown that arousal increases selectivity for a variety of stimulus types. For example, a previous perception study found that a tone conditioned to predict shock enhanced FFA activity in young adults when viewing salient face images and suppressed PPA activity when viewing non-salient scene images (Lee et al., 2013). A later memory study found that threat of punishment increased PPA activity associated with encoding goal-relevant scenes and suppressed lateral occipital cortex activity associated with encoding goal-irrelevant objects in young adults (Clewett et al., 2018). The age difference we observed in the arousal-by-salience interaction also corroborates another recent perception study which showed that a tone conditioned to predict shock increased neural gain in the PPA when viewing salient scenes in young but not older adults (Lee et al., 2018). Our findings thus show that the effect of arousal on selectivity is not limited to stimulus type (faces, scenes, bodies, objects), experimental design (memory or perception), or experiment specific features like task-related demands. Rather, arousal seems to more generally increase the brain’s selectivity in young but not older adults.

However, the prior evidence that arousal enhances selective processing in the PPA (Clewett et al., 2018; Lee et al., 2013, 2018) begs the question of why, in the current study, arousal modulated selectivity in young adults’ EBA and not PPA. One possibility is that the biological relevance of human bodies makes them more attention grabbing than scenes. Biologically relevant emotional stimuli (e.g., images relevant to survival or reproduction) have shown to modulate cognitive processing more automatically than socially relevant emotional stimuli (e.g., images related to social adaptation), the latter of which evoke elaborative processing in the frontal lobe (Sakaki et al., 2012). This implies that biologically relevant stimuli may be more likely to capture attention than non-biologically relevant stimuli. Consistent with this notion, studies have shown that body and face pictures are more rapidly processed during visual search (Ro et al., 2007) and more likely to be detected during an inattentional blindness task (Downing et al., 2004) than other objects (e.g., clothes, appliances, food, plants). Body stimuli also hold attention longer than other types of stimuli (Ro et al., 2007), which could explain why young adults were slower to detect probes appearing behind salient versus non-salient body images during our encoding task. Together, this evidence suggests that body images have an advantage in visual processing and, according to our results, increases in arousal may have an additive effect on this advantage.

It is important to highlight that the age difference in the arousal-by-salience interaction in the EBA cannot be attributed to age-related changes in category-selective processing. Despite evidence that neural processes become less distinct with age (Carp et al., 2011; Li et al., 2001), we observed category-selective responses in young and older adults. According to our whole-brain analysis, both age groups showed greater activation in category-selective occipitotemporal regions, including the EBA and PPA, when they were viewing salient versus non-salient images of bodies and scenes, respectively. Our results thus suggest that selective processing in these regions is intact in older adults (see also Lee et al., 2018).

Our findings also imply that the age difference in arousal-induced selectivity cannot be attributed to age differences in the arousal response. Relative to non-arousing sounds, both young and older adults in the current study showed higher subjective arousal ratings in response to arousing sounds. Measures of LC function were also similar across age groups as both pupil dilation and LC ROI activity were increased on arousing compared to non-arousing trials. The neuroimaging results further showed that, for both groups, arousing trials increased activation in brain regions consistent with the ventral attention network in the ventral frontal cortex and the rTPJ (Corbetta et al., 2008; Vossel et al., 2014) as well as the frontal orbital cortex, which is thought to play a role in driving phasic LC activation (Aston-Jones & Cohen, 2005). However, the arousal manipulation selectively modulated activity in the rTPJ as a function of salience for young adults but not for older adults. These results indicate that it is not likely the *response* to arousal that changes with age, but the efficiency of arousal in coordinating attention-related processes in the ventral attention network to amplify selective processing.

Finally, there were no age differences in LC contrast ratios. This aligns with previous findings in healthy adults (Hämmerer et al., 2018), as studies have shown that LC MRI contrast has a quadratic relationship with age, peaking at 60 years, remaining stable until ∼80 years, and gradually decreasing in the ninth decade (Liu et al., 2020; Shibata et al., 2006). A novel component of this study was the ability to test associations between LC MRI contrast and LC function. Given that LC MRI contrast is thought to reflect LC structural integrity (Liu et al., 2017), we expected to see correspondence of this measure with either LC ROI activity or pupil dilation, our independent measure of LC function (indeed, pupil dilation and LC ROI activity were positively correlated in young adults). However, we found no meaningful associations between LC structural MRI contrast and function. Nevertheless, LC contrast ratios were associated with better memory selectivity on neutral trials in young adults, consistent with prior findings of a positive relationship between young adults’ memory and LC MRI contrast (Clewett et al. 2018; Dahl et al. 2019). Limitations imposed by our small sample size could explain why anticipated associations were not present. Another contributing factor could be differences in imaging protocols used for structural LC assessment as well as the coordinates used for functional localization of the LC. Additional research is therefore needed to clarify the relationship between LC structure and LC function in young and older adults.

### 4.1 Conclusions

In conclusion, our findings indicate a dissociation in the way that arousal modulates selectivity in young and older adults’ behavior, category-selective processing, and ventral attention network activity. During increased emotional arousal, young adults are better able to process salient information, whereas older adults’ attention becomes less selective. Although older adults tend to show poorer selective attention and increased susceptibility to distraction than young adults (Hasher & Zacks, 1988), our results suggest that arousal may further exacerbate this vulnerability. In turn, older adults may be less able to focus during arousing situations, whether evoked by stimuli associated with an aversive event (Lee et al., 2018), a threat of punishment (Durbin et al., 2018), or from hearing an emotionally arousing sound. This could have detrimental consequences for older adults’ performance in high-stake moments.

## Acknowledgements

This research was supported by a BrightFocus Postdoctoral Fellowship (A2018449F) awarded to SG with partial support from Alzheimer’s Los Angeles. BK was supported by a National Institutes of Health grant (F32AG057162). SB was supported by National Science Foundation grant DGE-1842487 and National Institutes of Health grant T32AG000037. MM was supported by a National Institutes of Health grant (R01AG025340). We thank Christine Cho, Taylor Shigezawa, and Ilana Cohen for assistance with neuroimaging preprocessing and manual LC delineation as well as Katherin Martin for her help with MRI acquisition.

## Data Statement

Behavioral data will be available at https://osf.io/4276j/ and MRI data at OpenNeuro.org.

## Supplementary Material

### 1.0 Supplementary Methods

#### 1.1 Automated LC delineation

LC delineation was performed using a semi-automated procedure which has been previously described (Dahl et al. 2019). The TSE and MPRAGE scans were resampled to twice their native resolution, after which MPRAGE scans were pooled to create a whole-brain template. Resampled TSE scans were coregistered to whole-brain template-coregistered MPRAGE scans, and resulting scans were pooled to generate an TSE template. Following coregistration of the TSE template to the MPRAGE template and the MPRAGE template to MNI 0.5mm linear space, transformations from previous steps were used to warp resampled TSE scans and the TSE template to MNI 0.5mm linear space.

The ANTs routines and parameters used for LC delineation were the same as those described by Dahl et al. (2019) except for the following deviations. First, resampling of scans was performed with the ANTs ResampleImage routine. Second, template building was performed using the antsMultivariateTemplateConstruction.sh routine. Third, for construction of the initial (as opposed to the full) whole-brain template, we used a subset of 27 MPRAGE scans (16 from younger adults, 11 from older adults) all of which had `qoffset_x`, `qoffset_y`, and `qoffset_z` values within 1 standard deviation of the mean across all scans. This resulted in an initial template with high spatial alignment which was then used for alignment of all scans during construction of the whole-brain template. Finally, TSE scans and the TSE template were warped to MNI 0.5mm linear space, rather than whole-brain template space, for the purpose of comparing locations of hyperintensities on TSE scans and the TSE template with available LC maps (Dahl et al. 2019; Ye et al. 2021).

### 2.0 Supplementary Results

#### 2.1 Comparison of manual and automatic calculation of LC intensity

To validate the automated LC delineation approach, we compared the peak LC intensities derived from this method with those derived from a manual LC anatomical tracing procedure that has been previously described (Clewett et al. 2018; Clewett et al. 2016; Shibata et al. 2006). For the manual approach, left and right LC ROIs were hand drawn by two blinded raters on each participant’s native-resolution TSE T1-weighted image using FSLeyes image viewer. In the axial plane, raters first identified the slice where LC signal intensities were most apparent near the floor of the fourth ventricle. A 1×1mm ROI was then drawn on the voxel with peak intensity in each hemisphere. To measure reference intensity, a 10×10mm ROI was drawn on the dorsal pontine tegmentum, placed six voxels above and equidistant between the left and right LC ROIs in the axial plane. Intensity values were then extracted from the three ROIs and LC contrast ratios were calculated using the same LC contrast equation described in Section 2.6.3. Intraclass correlations coefficients between raters indicated high interrater reliability for LC peak intensities in the left (ICC = .94, *p* < .001, 95% CI [.899, .963]) and right hemispheres (ICC = .84, *p* < .001, 95% CI [.747, .903]). With high accordance established, peak intensities were averaged across raters for each hemisphere. Comparisons of the manually and automatically derived peak LC intensities revealed high correspondence for the left (ICC = .93, *p* < .001, 95% CI [.879, .955]) and right hemisphere (ICC = .90, *p* < .001, 95% CI [.837, .939]).

## Footnotes

1 As mentioned in the methods, one older adult’s MRI data was not saved on the scanner due to a technical error. Although we could not include their imaging data, we included their behavioral data in our analyses.

2 One older adult’s recognition data was lost due to a technical error.

## References

Aston-Jones, G., & Cohen, J. D. (2005). An integrative theory of locus coeruleus-norepinephrine function: Adaptive gain and optimal performance. Annual Review of Neuroscience, 28, 403– 450. https://doi.org/10.1146/annurev.neuro.28.061604.135709

Avants, B. B., Tustison, N., & Song, G. (2009). Advanced Normalization Tools (ANTS). The Insight Journal, 2(365), 1–35.

Beckmann, C. F., Jenkinson, M., & Smith, S. M. (2003). General multilevel linear modeling for group analysis in FMRI. NeuroImage, 20, 1052–1063. https://doi.org/10.1016/S1053-8119(03)00435-X

Betts, M. J., Cardenas-blanco, A., Kanowski, M., Jessen, F., & Düzel, E. (2017). In vivo MRI assessment of the human locus coeruleus along its rostrocaudal extent in young and older adults. NeuroImage, 163, 150–159. https://doi.org/10.1016/j.neuroimage.2017.09.042

Bowling, J. T., Friston, K. J., & Hopfinger, J. B. (2020). Top-down versus bottom-up attention differentially modulate frontal–parietal connectivity. Human Brain Mapping, 41(4), 928– 942. https://doi.org/10.1002/hbm.24850

Braak, H., & Del Tredici, K. (2011). The pathological process underlying Alzheimer’s disease in individuals under thirty. Acta Neuropathologica, 121(2), 171–181. https://doi.org/10.1007/s00401-010-0789-4

Bradley, M. M., & Lang, P. J. (2007). The International Affective Digitized Sounds (2nd Edition; IADS-2): Affective ratings of sounds and instruction manual (Technical report B-3). University of Florida, Gainesville, Fl.

Campbell, K. L., Grady, C. L., Ng, C., & Hasher, L. (2012). Age differences in the frontoparietal cognitive control network: Implications for distractibility. Neuropsychologia, 50(9), 2212– 2223. https://doi.org/10.1016/j.neuropsychologia.2012.05.025

Carp, J., Park, J., Polk, T. A., & Park, D. C. (2011). Age differences in neural distinctiveness revealed by multi-voxel pattern analysis. NeuroImage, 56(2), 736–743. https://doi.org/10.1016/j.neuroimage.2010.04.267

Clewett, D. V., Huang, R., Velasco, R., Lee, T. H., & Mather, M. (2018). Locus coeruleus activity strengthens prioritized memories under arousal. Journal of Neuroscience, 38(6), 1558–1574. https://doi.org/10.1523/JNEUROSCI.2097-17.2017

Corbetta, M., Patel, G., & Shulman, G. L. (2008). The reorienting system of the human brain: From environment to theory of mind. Neuron, 58(3), 306–324. https://doi.org/10.1016/j.neuron.2008.04.017

Corbetta, M., & Shulman, G. L. (2002). Control of goal-directed and stimulus-driven attention in the brain. Nature Reviews. Neuroscience, 3(3), 201–215. https://doi.org/10.1038/nrn755

Dahl, M. J., Mather, M., Düzel, S., Bodammer, N. C., Lindenberger, U., Kühn, S., & Werkle-Bergner, M. (2019). Rostral locus coeruleus integrity is associated with better memory performance in older adults. Nature Human Behaviour, 3(11), 1203–1214. https://doi.org/10.1038/s41562-019-0715-2

Dahl, M. J., Mather, M., Werkle-Bergner, M., Kennedy, B. L., Qiao, Y., Shi, Y., & Ringman, J. M. (2020). Lower MRI-indexed locus coeruleus integrity in autosomal-dominant Alzheimer’s disease. In Alzheimer’s & Dementia (Vol. 16, Issue S5). https://doi.org/10.1002/alz.047676

Downing, P. E., Bray, D., Rogers, J., & Childs, C. (2004). Bodies capture attention when nothing is expected. Cognition, 93(1), 27–38. https://doi.org/10.1016/j.cognition.2003.10.010

Downing, P. E., Chan, A. W. Y., Peelen, M. V., Dodds, C. M., & Kanwisher, N. (2006). Domain specificity in visual cortex. Cerebral Cortex, 16(10), 1453–1461. https://doi.org/10.1093/cercor/bhj086

Downing, P. E., Jiang, Y., Shuman, M., & Kanwisher, N. (2001). A cortical area selective for visual processing of the human body. Science, 293(5539), 2470–2473. https://doi.org/10.1126/science.1063414\r293/5539/2470 [pii]

Durbin, K. A., Clewett, D., Huang, R., & Mather, M. (2018). Age differences in selective memory of goal-relevant stimuli under threat. Emotion, 18(6), 906–911. https://doi.org/10.1037/emo0000398

Elman, J. A., Puckett, O. K., Beck, A., Fennema-Notestine, C., Cross, L. K., Dale, A. M., Eglit, G. M. L., Eyler, L. T., Gillespie, N. A., Granholm, E. L., Gustavson, D. E., Hagler, D. J., Jr, Hatton, S. N., Hauger, R., Jak, A. J., Logue, M. W., McEvoy, L. K., McKenzie, R. E., Neale, M. C., … Kremen, W. S. (2021). MRI-assessed locus coeruleus integrity is heritable and associated with multiple cognitive domains, mild cognitive impairment, and daytime dysfunction. Alzheimer’s & Dementia: The Journal of the Alzheimer’s Association, 17(6), 1017–1025. https://doi.org/10.1002/alz.12261

Epstein, R., & Kanwisher, N. (1998). The parahippocampal place area: A cortical representation of the local visual environment. NeuroImage, 392, 6–9. https://doi.org/10.1016/s1053-8119(18)31174-1

Fox, M. D., Corbetta, M., Snyder, A. Z., Vincent, J. L., & Raichle, M. E. (2006). Spontaneous neuronal activity distinguishes human dorsal and ventral attention systems. Proceedings of the National Academy of Sciences of the United States of America, 103(26), 10046–10051. https://doi.org/10.1073/pnas.0604187103

Gallant, S. N., Durbin, K. A., & Mather, M. (2020). Age differences in vulnerability to distraction under arousal. Psychology and Aging, 35(5), 780–791. https://doi.org/10.1037/pag0000426

Griffanti, L., Douaud, G., Bijsterbosch, J., Evangelisti, S., Alfaro-Almagro, F., Glasser, M. F., Duff, E. P., Fitzgibbon, S., Westphal, R., Carone, D., Beckmann, C. F., & Smith, S. M. (2017). Hand classification of fMRI ICA noise components. NeuroImage, 154, 188–205. https://doi.org/10.1016/j.neuroimage.2016.12.036

Hämmerer, D., Callaghan, M. F., Hopkins, A., Kosciessa, J., Betts, M., Cardenas-Blanco, A., Kanowski, M., Weiskopf, N., Dayan, P., Dolan, R. J., Düzel, E., Kanowski, M., Kosciessa, J., Betts, M., Callaghan, M. F., Düzel, E., Hopkins, A., Weiskopf, N., & Hämmerer, D. (2018). Locus coeruleus integrity in old age is selectively related to memories linked with salient negative events. Proceedings of the National Academy of Sciences of the United States of America, 115(9), 2228–2233. https://doi.org/10.1073/pnas.1712268115

Hasher, L., & Zacks, R. T. (1988). Working memory, comprehension, and aging: A review and a new view. Psychology of Learning and Motivation, 22, 193–225. https://doi.org/10.1016/S0079-7421(08)60041-9

Jacobs, H. I. L., Priovoulos, N., Poser, B. A., Pagen, L. H. G., Ivanov, D., Verhey, F. R. J., & Uludağ, K. (2020). Dynamic behavior of the locus coeruleus during arousal-related memory processing in a multi-modal 7T fMRI paradigm. eLife, 9, 1–30. https://doi.org/10.7554/eLife.52059

Jenkinson, M., Bannister, P., Brady, M., & Smith, S. (2002). Improved optimization for the robust and accurate linear registration and motion correction of brain images. NeuroImage, 17, 825–841. https://doi.org/10.1006/nimg.2002.1132

Kahnt, T., & Tobler, P. N. (2013). Salience signals in the right temporoparietal junction facilitate value-based decisions. Journal of Neuroscience, 33(3), 863–869. https://doi.org/10.1523/JNEUROSCI.3531-12.2013

Kaiser, D., Strnad, L., Seidl, K. N., Kastner, S., & Peelen, M. V. (2014). Whole person-evoked fMRI activity patterns in human fusiform gyrus are accurately modeled by a linear combination of face-and body-evoked activity patterns. 82–90. https://doi.org/10.1152/jn.00371.2013

Kennedy, B. L., & Mather, M. (2019). Neural mechanisms underlying age-related changes in attentional selectivity. In G. R. Samanez-Larkin (Ed.), The aging brain: Functional adaptation across adulthood (pp. 45–72). American Psychological Association. https://doi.org/10.1037/0000143-003

Keren, N. I., Lozar, C. T., Harris, K. C., Morgan, P. S., & Eckert, M. A. (2009). In vivo mapping of the human locus coeruleus. NeuroImage, 47(4), 1261–1267. https://doi.org/10.1016/j.neuroimage.2009.06.012

Klein, R. M. (2000). Inhibition of return. Trends in Cognitive Sciences, 4(4), 138–147. https://doi.org/10.1016/S1364-6613(00)01452-2

Kuznetsova, A., Brockhoff, P. B., & Christensen, R. H. B. (2017). lmerTest package: Tests in linear mixed effects models. Journal of Statistical Software, 82(13). https://doi.org/10.18637/jss.v082.i13

Lee, T. H., Greening, S. G., Ueno, T., Clewett, D., Ponzio, A., Sakaki, M., & Mather, M. (2018). Arousal increases neural gain via the locus coeruleus-noradrenaline system in younger adults but not in older adults. Nature Human Behaviour, 2(5), 356–366. https://doi.org/10.1038/s41562-018-0344-1

Lee, T. H., Sakaki, M., Cheng, R., Velasco, R., & Mather, M. (2013). Emotional arousal amplifies the effects of biased competition in the brain. Social Cognitive and Affective Neuroscience, 9(12), 2067–2077. https://doi.org/10.1093/scan/nsu015

Li, S. C., Lindenberger, U., & Sikström, S. (2001). Aging cognition: from neuromodulation to representation. Trends in Cognitive Sciences, 5(11), 479–486. https://doi.org/10.1016/s1364-6613(00)01769-1

Liu, K. Y., Kievit, R. A., Tsvetanov, K. A., Betts, M. J., Düzel, E., Rowe, J. B., Tyler, L. K., Brayne, C., Bullmore, E. T., Calder, A. C., Cusack, R., Dalgleish, T., Duncan, J., Henson, R. N., Matthews, F. E., Marslen-Wilson, W. D., Rowe, J. B., Shafto, M. A., Campbell, K., … Hämmerer, D. (2020). Noradrenergic-dependent functions are associated with age-related locus coeruleus signal intensity differences. Nature Communications, 11(1), 1–9. https://doi.org/10.1038/s41467-020-15410-w

Liu, K. Y., Marijatta, F., Hämmerer, D., Acosta-Cabronero, J., Düzel, E., & Howard, R. J. (2017). Magnetic resonance imaging of the human locus coeruleus: A systematic review. Neuroscience and Biobehavioral Reviews, 83(September), 325–355. https://doi.org/10.1016/j.neubiorev.2017.10.023

Lovibond, P. F., & Lovibond, S. H. (1995). The structure of negative emotional states: Comparison of the Depression Anxiety Stress Scales (DASS) with the Beck Depression and Anxiety Inventories. Behaviour Research and Therapy, 33(3), 335–343. https://doi.org/10.1016/0005-7967(94)00075-U

Mars, R. B., Sallet, J., Schüffelgen, U., Jbabdi, S., Toni, I., & Rushworth, M. F. S. (2012). Connectivity-based subdivisions of the human right “temporoparietal junction area”: Evidence for different areas participating in different cortical networks. Cerebral Cortex, 22(8), 1894–1903. https://doi.org/10.1093/cercor/bhr268

Mather, M. (2020). The locus coeruleus-norepinephrine system role in cognition and how it changes with aging. In D. Poeppel, G. Mangun, & M. Gazzaniga (Eds.), The Cognitive Neurosciences (6th ed., pp. 91–104). The MIT Press. https://www.ncbi.nlm.nih.gov/pubmed/25246403

Mather, M., Clewett, D., Sakaki, M., & Harley, C. W. (2016). Norepinephrine ignites local hotspots of neuronal excitation: How arousal amplifies selectivity in perception and memory. The Behavioral and Brain Sciences, 39(200). https://doi.org/10.1017/S0140525X15000667

Mather, M., & Harley, C. W. (2016). The locus coeruleus: Essential for maintaining cognitive function and the aging brain. Trends in Cognitive Sciences, 20(3), 214–226. https://doi.org/10.1016/j.tics.2016.01.001

Mather, M., & Sutherland, M. R. (2011). Arousal-biased competition in perception and memory. Perspectives on Psychological Science: A Journal of the Association for Psychological Science, 6(2), 114–133. https://doi.org/10.1177/1745691611400234

Mathôt, S., van der Linden, L., Grainger, J., & Vitu, F. (2013). The pupillary light response reveals the focus of covert visual attention. PloS One, 8(10). https://doi.org/10.1371/journal.pone.0078168

Meteyard, L., & Davies, R. A. I. (2020). Best practice guidance for linear mixed-effects models in psychological science. Journal of Memory and Language, 112(March 2019), 104092. https://doi.org/10.1016/j.jml.2020.104092

Nasreddine, Z. S., Phillips, N. A., Bédirian, V., Charbonneau, S., Whitehead, V., Collin, I., Cummings, J. L., & Chertkow, H. (2005). The Montreal Cognitive Assessment, MoCA: A brief screening tool for mild cognitive impairment. Journal of the American Geriatrics Society, 53(4), 695–699. https://doi.org/10.1111/j.1532-5415.2005.53221.x

Nieuwenhuis, S., Aston-Jones, G., & Cohen, J. D. (2005). Decision making, the P3, and the locus coeruleus-norepinephrine system. Psychological Bulletin, 131(4), 510–532. https://doi.org/10.1037/0033-2909.131.4.510

Orlov, T., Makin, T. R., & Zohary, E. (2010). Topographic Representation of the Human Body in the Occipitotemporal Cortex. Neuron, 68(3), 586–600. https://doi.org/10.1016/j.neuron.2010.09.032

Posner, M. I., & Cohen, Y. (1984). Components of visual orienting. In H. Bouma & D. G. Bouwhuis (Eds.), Attention and performance X: Control of language processes (pp. 531– 556). Lawrence Erlbaum.

Ptak, R. (2012). The frontoparietal attention network of the human brain: Action, saliency, and a priority map of the environment. The Neuroscientist: A Review Journal Bringing Neurobiology, Neurology and Psychiatry, 18(5), 502–515. https://doi.org/10.1177/1073858411409051

Robertson, I. H. (2013). A noradrenergic theory of cognitive reserve: Implications for Alzheimer’s disease. Neurobiology of Aging, 34(1), 298–308. https://doi.org/10.1016/j.neurobiolaging.2012.05.019

Ro, T., Friggel, A., & Lavie, N. (2007). Attentional biases for faces and body parts. Visual Cognition, 15(3), 322–348. https://doi.org/10.1080/13506280600590434

Sakaki, M., Niki, K., & Mather, M. (2012). Beyond arousal and valence: the importance of the biological versus social relevance of emotional stimuli. Cognitive, Affective & Behavioral Neuroscience, 12(1), 115–139. https://doi.org/10.3758/s13415-011-0062-x

Samuels, E., & Szabadi, E. (2008). Functional neuroanatomy of the noradrenergic locus coeruleus: Its roles in the regulation of arousal and autonomic function part I: Principles of functional organisation. Current Neuropharmacology, 6(3), 235–253. https://doi.org/10.2174/157015908785777229

Sara, S. J. (2009). The locus coeruleus and noradrenergic modulation of cognition. Nature Reviews. Neuroscience, 10(3), 211–223. https://doi.org/10.1038/nrn2573

Sara, S. J., & Bouret, S. (2012). Orienting and reorienting: The locus coeruleus mediates cognition through arousal. Neuron, 76(1), 130–141. https://doi.org/10.1016/j.neuron.2012.09.011

Shibata, E., Sasaki, M., Tohyama, K., Kanbara, Y., Otsuka, K., Ehara, S., & Sakai, A. (2006). Age-related changes in locus ceruleus on neuromelanin magnetic resonance imaging at 3 Tesla. Magnetic Resonance in Medical Sciences: MRMS: An Official Journal of Japan Society of Magnetic Resonance in Medicine, 5(4), 197–200. https://doi.org/10.2463/mrms.5.197

Shipley, W. C. (1940). A self-administering scale for measuring intellectual impairment and deterioration. The Journal of Psychology, 9(2), 371–377. https://doi.org/10.1080/00223980.1940.9917704

Shomstein, S. (2012). Cognitive functions of the posterior parietal cortex: Top-down and bottom-up attentional control. Frontiers in Integrative Neuroscience, 6(Article 38), 1–7. https://doi.org/10.3389/fnint.2012.00038

Strange, B. A., & Dolan, R. J. (2007). ?-adrenergic modulation of oddball responses in humans. Behavioral and Brain Functions: BBF, 3(29), 1–5. https://doi.org/10.1186/1744-9081-3-29

Taylor, J. C., Wiggett, A. J., & Downing, P. E. (2007). Functional MRI analysis of body and body part representations in the extrastriate and fusiform body areas. Journal of Neurophysiology, 98(3), 1626–1633. https://doi.org/10.1152/jn.00012.2007

Vossel, S., Geng, J. J., & Fink, G. R. (2014). Dorsal and ventral attention systems: Distinct neural circuits but collaborative roles. The Neuroscientist: A Review Journal Bringing Neurobiology, Neurology and Psychiatry, 20(2), 150–159. https://doi.org/10.1177/1073858413494269

Weinshenker, D. (2018). Long road to ruin: Noradrenergic dysfunction in neurodegenerative disease. Trends in Neurosciences, 41(4), 211–223. https://doi.org/10.1016/j.tins.2018.01.010

Willenbockel, V., Sadr, J., Fiset, D., Horne, G. O., Gosselin, F., & Tanaka, J. W. (2010). Controlling low-level image properties: The SHINE toolbox. Behavior Research Methods, 42(3), 671–684. https://doi.org/10.3758/BRM.42.3.671

Xiao, J., Hays, J., Ehinger, K. A., Oliva, A., & Torralba, A. (2010). SUN database: Large-scale scene recognition from abbey to zoo. Proceedings of the IEEE Computer Society Conference on Computer Vision and Pattern Recognition, 3485–3492. https://doi.org/10.1109/CVPR.2010.5539970

Ye, R., Rua, C., O’Callaghan, C., Jones, P. S., Hezemans, F. H., Kaalund, S. S., Tsvetanov, K. A., Rodgers, C. T., Williams, G., Passamonti, L., & Rowe, J. B. (2021). An in vivo probabilistic atlas of the human locus coeruleus at ultra-high field. NeuroImage, 225, 117487. https://doi.org/10.1016/j.neuroimage.2020.117487

## Supplementary References

Avants, B. B., Tustison, N., & Song, G. (2009). Advanced normalization tools (ANTS). Insight J, 2(365), 1–35.

Dahl, M. J., Mather, M., Werkle-Bergner, M., Kennedy, B. L., Guzman, S., Hurth, K., Miller, C. A., Qiao, Y., Shi, Y., Chui, H. C., & Ringman, J. M. (2020). Locus coeruleus integrity is related to tau burden and memory loss in autosomal-dominant Alzheimer’s disease [Preprint]. Neurology. https://doi.org/10.1101/2020.11.16.20232561

Keren, N., Lozar, C. T., Harris, K. C., Morgan, P. S., & Eckert, M. A. (2009). In-Vivo Mapping of the Human Locus Coeruleus. NeuroImage, 47(4), 1261–1267. https://doi.org/10.1016/j.neuroimage.2009.06.012

Liu, K. Y., Marijatta, F., Hämmerer, D., Acosta-Cabronero, J., Düzel, E., & Howard, R. J. (2017). Magnetic resonance imaging of the human locus coeruleus: A systematic review. Neuroscience & Biobehavioral Reviews, 83, 325–355. https://doi.org/10.1016/j.neubiorev.2017.10.023

Ye, R., Rua, C., O’Callaghan, C., Jones, P. S., Hezemans, F. H., Kaalund, S. S., Tsvetanov, K. A., Rodgers, C. T., Williams, G., Passamonti, L., & Rowe, J. B. (2021). An in vivo probabilistic atlas of the human locus coeruleus at ultra-high field. NeuroImage, 225, 117487. https://doi.org/10.1016/j.neuroimage.2020.1174

